# CCPLS reveals cell-type-specific spatial dependence of transcriptomes in single cells

**DOI:** 10.1101/2022.01.12.476034

**Authors:** Takaho Tsuchiya, Hiroki Hori, Haruka Ozaki

**Affiliations:** Bioinformatics Laboratory, Faculty of Medicine, University of Tsukuba. Tsukuba 1-1-1 Tennodai, Tsukuba, Ibaraki 305-8577, Japan; Center for Artificial Intelligence Research, University of Tsukuba, 1-1-1 Tennodai, Tsukuba, Ibaraki 305-8577, Japan; Doctoral Program in Medical Sciences, Graduate School of Comprehensive Human Sciences, University of Tsukuba, 1-1-1 Tennodai, Tsukuba, Ibaraki 305-8577, Japan

## Abstract

**Motivation:** Cell-cell communications regulate internal cellular states, e.g., gene expression and cell functions, and play pivotal roles in normal development and disease states. Furthermore, single-cell RNA sequencing methods have revealed cell-to-cell expression variability of highly variable genes (HVGs), which is also crucial. Nevertheless, the regulation on cell-to-cell expression variability of HVGs via cell-cell communications is still largely unexplored. The recent advent of spatial transcriptome methods has linked gene expression profiles to the spatial context of single cells, which has provided opportunities to reveal those regulations. The existing computational methods extract genes with expression levels influenced by neighboring cell types. However, limitations remain in the quantitativeness and interpretability: they neither focus on HVGs nor consider the effects of multiple neighboring cell types.

**Results:** Here, we propose CCPLS (Cell-Cell communications analysis by Partial Least Square regression modeling), which is a statistical framework for identifying cell-cell communications as the effects of multiple neighboring cell types on cell-to-cell expression variability of HVGs, based on the spatial transcriptome data. For each cell type, CCPLS performs PLS regression modeling and reports coefficients as the quantitative index of the cell-cell communications. Evaluation using simulated data showed our method accurately estimated the effects of multiple neighboring cell types on HVGs. Furthermore, applications to the two real datasets demonstrate that CCPLS can extract biologically interpretable insights from the inferred cell-cell communications.

**Availability:** The R package is available at https://github.com/bioinfo-tsukuba/CCPLS. The data are available at https://github.com/bioinfo-tsukuba/CCPLS_paper.

**Supplementary information:** Supplementary data are available at Bioinformatics online.

## 1 Introduction

Cell-cell communications regulate internal cellular states, e.g., gene expression and cell functions, and play pivotal roles in normal development and disease states (Sharpe and Pauken, 2018; Snijder and Pelkmans, 2011; Pelkmans, 2012). For example, in tumor progression, programmed cell death ligand 1 (PD-L1) is expressed in a tumor cell that binds to the programmed cell death protein 1 (PD-1) of T cells, thus regulating T cell gene expression and weakening the anti-tumor growth response (Shimizu et al., 2020). The PD-1 pathway inhibitors block this cellcell communication and thus stop the growth of tumor cells (Sharpe and Pauken, 2018). Accordingly, it is essential to elucidate gene expression regulation via cell-cell communications in order to understand and control complex multicellular systems (Sharpe and Pauken, 2018; Snijder and Pelkmans, 2011; Pelkmans, 2012).

Single-cell RNA sequencing (scRNA-seq) methods have been used to study complex multicellular systems. These technologies have revealed cell-to-cell expression variability of highly variable genes (HVGs) even in the same cell type, which is crucial in the normal development and disease states (Ben-Moshe and Itzkovitz, 2019; Satija et al., 2015; Stuart et al., 2019). Nevertheless, the regulation on cell-to-cell expression variability of HVGs via cell-cell communications has not been well characterized (Ben-Moshe and Itzkovitz, 2019; Satija et al., 2015; Stuart et al., 2019). Instead of the expression of HVGs, the expression of ligand and receptor genes have been the focus of the computational tools to infer potential cell-cell communications using scRNA-seq data: these tools compare expression levels of ligand and receptor genes between cell-type pairs (Armingol *et al*., 2020; Hou *et al*., 2020; Nagai *et al*., 2021). However, such approaches have several limitations regarding incompleteness of knowledge on the ligand-receptor pairs, potential crosstalks among ligands and receptors, and unavailability of spatial contexts of cells (Armingol *et al*., 2020; Hou *et al*., 2020; Nagai *et al*., 2021). Moreover, the ligand-receptor expression approach cannot directly provide insights into the effects of cell-cell communications in most of the HVGs in the same cell type. It is desirable to focus on the regulation on cell-to-cell expression variability of HVGs via cell-cell communications.

As another remarkable aspect, intracellular gene expression regulation is a multiple-input and multiple-output (MIMO) system, and a quantitative understanding is of critical importance (Akimoto *et al*., 2013; Janes *et al*., 2005). The regulation via cell-cell communications can also be considered a MIMO system, in which the degree of neighboring cell-type existence regulates the intracellular gene expression values. As an example of the PD-1 pathway, antigen-presenting cells and the tumor cells cooperatively regulate the gene expression of T cells, and the balance of such regulation is crucial to cell fate decisions (Sharpe and Pauken, 2018). Besides cellcell communications via known molecular interactions, the cases in which the gene expression in a cell is affected by the spatial arrangement and combinations of the neighboring cells can be interpreted as MIMO systems (Colombo and Cattaneo, 2021; Haanen, 2017; Hui and Bhatia, 2007). It is essential to develop computational methods for estimating such a MIMO system of cell-cell communications by using spatial spatial contexts of cells.

The recent advent of spatial transcriptome methods has linked gene expression profiles to the spatial context of single cells, which has provided opportunities to reveal the regulation on cell-to-cell expression variability of HVGs via cell-cell communications (Cho *et al*., 2021; Eng *et al*., 2019; Hu *et al*., 2021b; Marx, 2021). Nevertheless, there is no computational methods based on spatial transcriptome data for estimating regulation on cell-to-cell expression variability of HVGs as a MIMO system of cell-cell communications (Arnol *et al*., 2019; Dries *et al*., 2021b; Hu *et al*., 2021a; Rao *et al*., 2021; Svensson *et al*., 2018; Tanevski *et al*., 2022; Velten *et al*., 2022; Zhu *et al*., 2021), which could limit quantitativeness and interpretability. For example, Giotto findICG extracts genes with expression levels that are influenced by the neighboring cell types; however, it neither considers the cooperation of neighboring cell types nor quantifies the degree of regulation with each neighboring cell type (Dries *et al*., 2021b). Other existing methods focus on the spatial gene expression variability itself or gene-gene relationships and do not focus on the effect of neighboring cell types on gene expression variability (Arnol *et al*., 2019; Dries *et al*., 2021b; Hu *et al*., 2021a; Rao *et al*., 2021; Svensson *et al*., 2018; Tanevski *et al*., 2022; Velten *et al*., 2022; Zhu *et al*., 2021). Moreover, these tools do not explicitly focus on HVGs. Thus, there is a need to develop methods to estimate the MIMO system of cell-cell communications that focus on HVGs.

Here, we propose CCPLS (Cell-Cell communications analysis by Partial Least Square regression modeling), which is a statistical framework for identifying the MIMO system of cell-cell communications, i.e., regulation on cell-to-cell expression variability of HVGs by multiple neighboring cell types, based on the spatial transcriptome data at a singlecell resolution. CCPLS performs PLS regression modeling for each cell type and reports the estimated coefficients as the quantitative index of the cell-cell communications from each neighboring cell type. Evaluation using simulated data showed our method accurately estimated the effects of multiple neighboring cell types on HVGs. Furthermore, by applying CCPLS to the two real datasets, we demonstrate CCPLS can be used to extract biologically interpretable insights from the inferred cell-cell communications.

## 2 Materials and Methods

### 2.1 Problem definition

Spatial transcriptome data at a single-cell resolution consist of a gene expression matrix, a coordinate matrix, and a cell-type label vector. Let *N* be the number of cells, *G* be the number of genes, and *M* be the number of cell types. The gene expression matrix 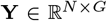 contains the expression value *y_i,g_* for each cell *i* (1 ≤ *i* ≤ *N*) and gene *g*(1 ≤ *g* ≤ *G*). The coordinate matrix 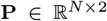 contains the twodimensional spatial coordinate of cells (*p*_*i*,1_, *p*_*i*,2_) (1 ≤ *i* ≤ *N*). The cell-type label vector **L** = {*l_i_*|*l_i_* ∈ {1,…,*M*}} contains the cell-type label *l_i_* of the M unique cell types. We assume the existence of highly variable genes (HVGs) specific to cell type *m* (1 ≤ *m* ≤ *M*), i.e., **h**^(*m*)^, which is estimated from the expression matrix **Y**^(*m*)^ containing cells i (1 ≤ *i* ≤ *N*^(*m*)^) of cell type *m* (Stuart *et al*., 2019; Hafemeister and Satija, 2019).

CCPLS does not estimate cell-cell communications via ligand-receptor interactions but estimates the effect on the gene expression of a cell type due to the arrangement and combination of neighboring cell types as a MIMO system. There are two key assumptions of CCPLS: (i) the cellcell communications are composed of the linear sum of the effects of neighboring cell types; (ii) the effects of neighboring cell types are the same within each cell type. Accordingly, CCPLS aims to extract HVGs **h**^(*m*)^ whose expression values are significantly explained by a linear sum of the effects by neighboring cell types *f* as follows:

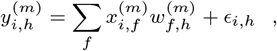

where *h* ∈ **h**^(*m*)^, 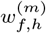 denotes coefficient, 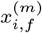 denotes neighboring cell-type score, and *ϵ_i,h_* is the residue term.

Simultaneously, CCPLS aims to estimate the coefficient 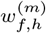 as the direction and degree of regulation via cell-cell communications by neighboring cell types *f* for each HVG *h* in a manner specific to cell type *m*.

### 2.2 CCPLS

#### 2.2.1 Methods overview

The core of CCPLS is PLS regression modeling performed by setting HVG expression values 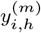 as responsive variables and neighboring cell-type scores 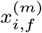 as explanatory variables for the each cell type *m*. After the statistical evaluation, CCPLS performs HVGs **h**^(*m*)^ clustering, which is reported with filtered coefficients as the direction and degree of regulation in the cell-cell communication. The input data for CCPLS are the expression matrix **Y**, the coordinate matrix **P**, and the cell-type label vector **L**. CCPLS consists of six main steps as described in sections 2.2.2–2.2.7, respectively.

Except for the calculation of neighboring cell-type scores, CCPLS is principally based on existing procedures. In other words, by calculating the neighboring cell-type scores, CCPLS realizes a multiple regression approach for estimating the MIMO system of cell-cell communications.

A schematic illustration of the MIMO system of cell-cell communications is shown in Figure 1a, while Figure 1b illustrates the workflow of CCPLS.

**Fig. 1.**
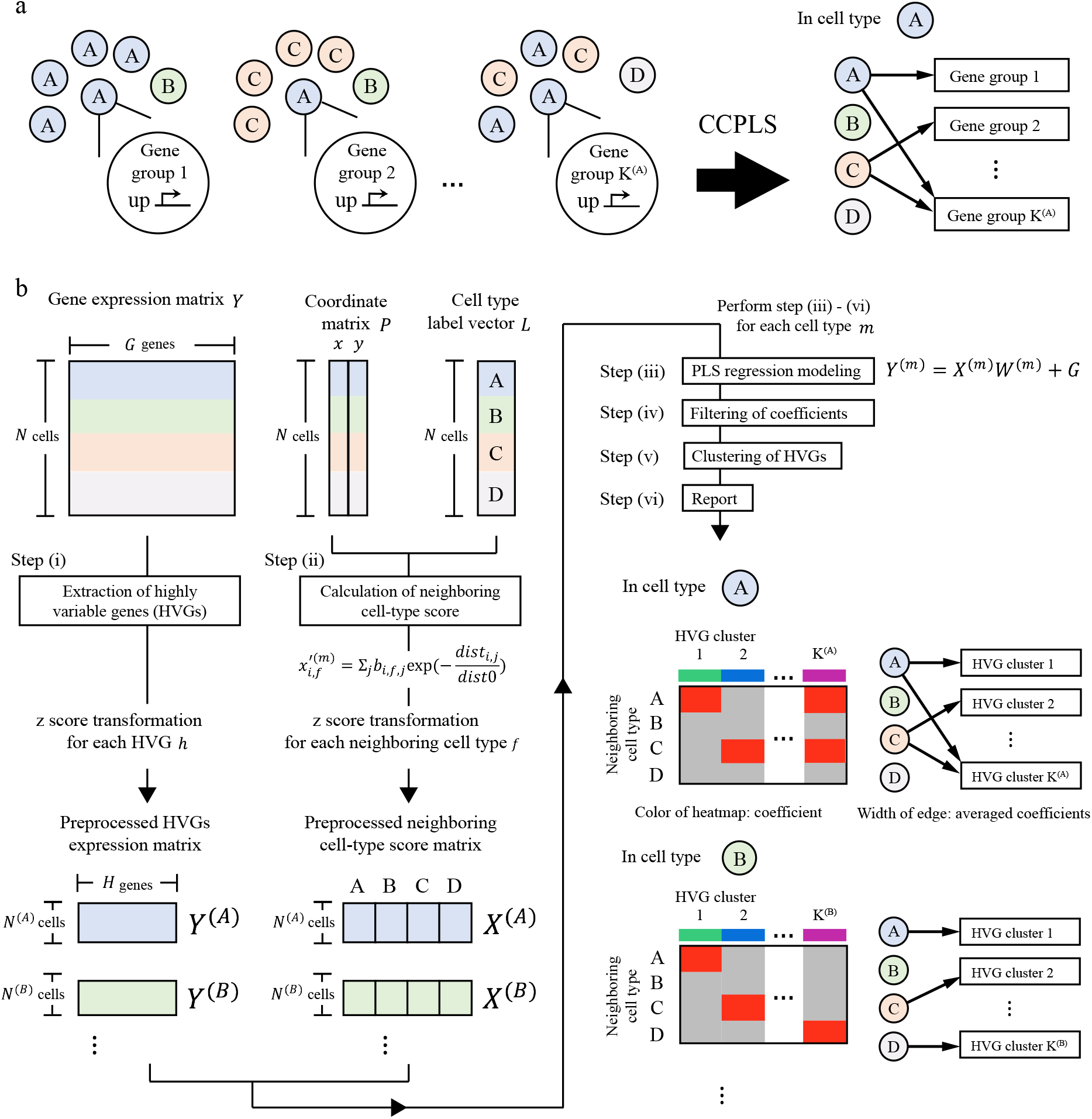
Overview of CCPLS. (a) Schematic illustration of the multiple-input and multiple-output (MIMO) system of cell-cell communications that CCPLS aims to identify. (b) Workflow of CCPLS.

#### 2.2.2 Step (i): Extraction of HVG

CCPLS divides the input expression matrix **Y** into each cell type **Y**′^(*m*)^, the expression matrix comprised of cells *i* for each unique cell type *m*. CCPLS filters out the genes with an expression value of 0 in all the cells within cell type *m* from **Y′**^(*m*)^, which is subsequently normalized and for each transformed gene z-score. Simultaneously, CCPLS extracts cell-type-*m*-specific HVG **h**^(*m*)^ and preprocessed HVG expression value 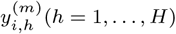 and matrix 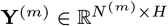 (Hafemeister and Satija, 2019; Stuart *et al*., 2019).

#### 2.2.3 Step (ii): Calculation of neighboring cell-type score

For each cell type *m*, CCPLS calculates the degree of neighboring cell-type existence 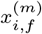, denoted as the neighboring cell-type score, by using the coordinate matrix **P** and cell-type label vector **L**. CCPLS calculates un-preprocessed neighboring cell-type score 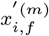 based on a function that decays with distance between two cells *i* and *j* as follows:

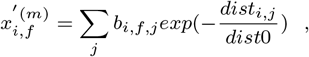

where the *b_i,f,j_* is the binary value indicating whether the cell type *l_j_* of cell *j* (1 ≤ *j* ≤ *N*, *j* ≠ *i*) belongs to cell type *f*, and the *dist_i,j_* is the Euclidean distance calculated from the coordinates of the two cells *i* and *j*. The value of *dist*_0_ is the constant set as the minimum value of the Euclidean distance of all the combinations of two cells in the coordinate matrix **P**. CCPLS performs z-score transformation on the un-preprocessed neighboring cell-type score 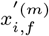 for each neighboring cell type f and calculates the preprocessed neighboring cell-type score matrix **X**^(*m*)^. As described here, CCPLS employs a Gaussian kernel similar to the previous studies (Arnol *et al*., 2019; Dang *et al*. (2020)).

#### 2.2.4 Step (iii): PLS regression modeling

For each cell type *m*, CCPLS performs PLS regression modeling (Höskuldsson, 1988). CCPLS employs PLS regression because its advantage is the capability of handling the “small N, large P” problem which is one of the characteristics of the spatial transcriptome data (Abdi, 2010; Nagasawa *et al*., 2021). The PLS regression model can be understood as two steps regression model (Akimoto *et al*., 2013; Höskuldsson, 1988).

The first step can be considered as consisting of the development of outer relations (**X** and **Y** metric individually). These data matrices are decomposed into latent variables plus a residue matrix. The sub-matrices can be represented as the product of the scores and the loadings which can be re-grouped in independent matrices as follows:

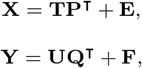

where 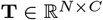 and 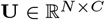 are the score matrices, and **P**^*L*×*C*^ and 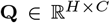 are the loading matrices, for the **X** and **Y** matrices, respectively. *C* is the component number for PLS regression. The matrices 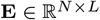 and 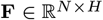 correspond to the residues associated with the PLS regression modeling.

The second step is a linear inner relation linking between **T** and **U**,

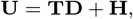

where 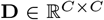 is the diagonal matrix and 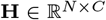 denotes the residue matrix. Here, PLS regression modeling yields by,

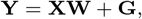

where 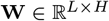 is the matrix of coefficients

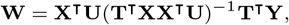

and 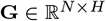 is the residue matrix.

For the each cell type *m*, CCPLS estimates this coefficient matrix **W**^(*m*)^ by the ***Y***^(*m*)^ and **X**^(*m*)^. **W**^(*m*)^ consists of the each coefficients 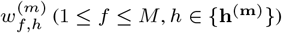, which are the direction and degree of regulation in the cell-cell communications which CCPLS aims to estimate. The each coefficient consists of each component *c* as the equation 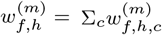.

CCPLS uses component number *C* that minimizes the Mean Squared Error (MSE) of 10-folds cross-validation (Bengio and Grandvalet, 2004).

#### 2.2.5 Step (vi): Filtering of coefficients

For each cell type *m*, CCPLS performs two-step filtering, a *t*-test of factor loadings, and a non-parametric test of the coefficients 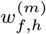 (Yamamoto *et al*., 2014). In each step, the *p*-values are false discovery rate (FDR)-adjusted by the Benjamini-Hochberg (BH) method, respectively (Benjamini and Hochberg, 1995). The first step filtered out coefficients in each component *w_f,h,c_* while remaining summed coefficients *w_f,h_* = ∑_*c*_*w_f,h,c_* with small values. Based on this tendency, CCPLS employed the two steps filtering. See text S1 in the Supplementary Information for the detailed procedures.

#### 2.2.6 Step (v): Clustering of HVGs

For each cell type *m*, CCPLS performs HVG clustering by the *k*-means method on the filtered coefficients (Yuan and Yang, 2019). The cluster number *k* is determined by the Silhouette method (Yuan and Yang, 2019). The minimum and the maximum integer values of *k* are set as 2 and 15, respectively. Before clustering, CCPLS filters out the HVGs whose filtered coefficients are all zero.

#### 2.2.7 Step (vi): Report

CCPLS outputs a heat map and bipartite graph. The heat map reports the filtered coefficients for each combination of HVGs *h* and neighboring cell types *f*. The bipartite graph visualizes the relationship between the HVG clusters and the neighboring cell types *f*, where the widths of edges are the averaged values of the filtered coefficients of each HVG cluster.

### 2.3 Datasets

#### 2.3.1 Simulated dataset

For the evaluation of CCPLS performance, we first simulated a dataset according to the following procedures. To realistically simulate the cell positions and ratio of cell types, our simulation experiments were based on the seqFISH+ real dataset (Eng *et al*., 2019).

We used the coordinate matrix **P** of the seqFISH+ real dataset. To simplify the cell types, we prepared a cell-type label vector **L** by substituting cell types A-D as the cell-type label vector of the seqFISH+ real dataset. We set four cell types as inputs and four HVG clusters as outputs, which contained 500 genes, respectively (Fig. 2a). HVG cluster 1 was a multiple-input case, while clusters 2-3 were single-input cases. HVG cluster 4 was not affected by neighboring cell types.

**Fig. 2.**
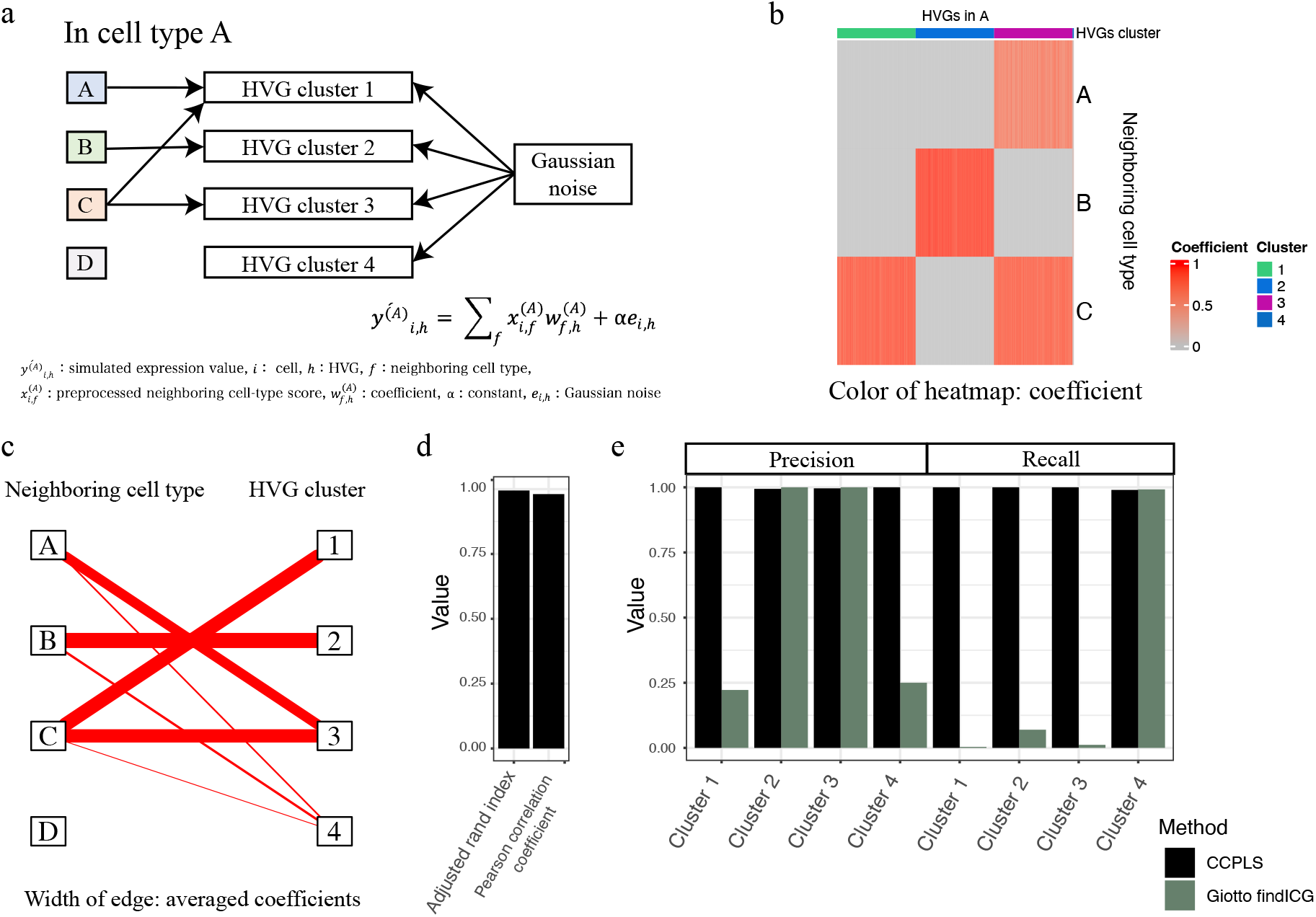
Evaluation using the simulated data. (a) Schematic illustration of the simulation settings. (b) Heat map generated by CCPLS. The color of the heat map indicates the coefficient. Rows and columns correspond to neighboring cell types and highly variable genes (HVGs), respectively. (c) Bipartite graph generated by CCPLS. The width of each edge indicates the average coefficients for each combination of HVG clusters and neighboring cell types. (d) Performance indexes and comparison with Giotto findICG. The calculated indexes are as follows: adjusted rand index, Pearson correlation coefficient, precision, and recall of each HVG cluster. Red and green indicate CCPLS and Giotto findICG results, respectively.

We simulated the preprocessed expression value of cell type A according to the equation 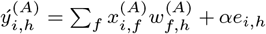. We prepared coefficient 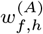 according to Table 1, in which the coefficient is *w_max_* or 0. We calculated the preprocessed neighboring cell-type score 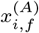 using the prepared coordinate matrix **P** and the cell-type label vector **L**. We simulated the term *e_i,h_* as Gaussian noise, with a mean of 0 and standard deviations of each respective HVG estimated by Mean-CV regression modeling of the seqFISH+ real dataset, which was then multiplied by the constant *α*, which were used for the evaluation of CCPLS performance (Fig. 2 and Fig. S1). Note that we did not perform experiments in count scale, which was originally un-recommended in the literature (Oshlack and Wakefield, 2009; Van Verk *et al*., 2013), and instead simulated scaled data.

**Table 1.**
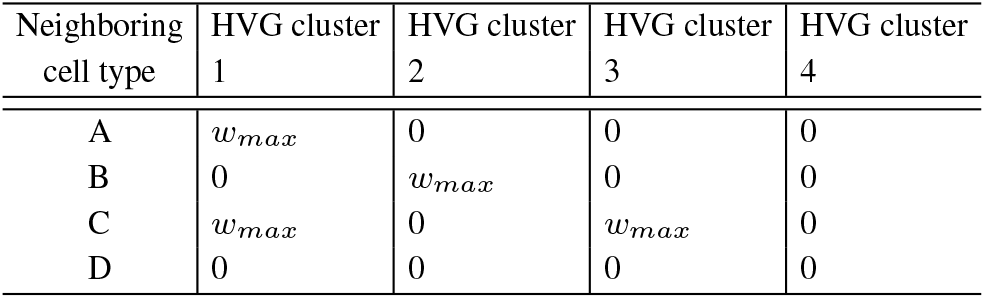
Coefficients 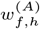 in the simulation experiments

| Neighboring cell type | HVG cluster 1 | HVG cluster 2 | HVG cluster 3 | HVG cluster 4 |
| --- | --- | --- | --- | --- |
| A | $w_{max}$ | 0 | 0 | 0 |
| B | 0 | $w_{max}$ | 0 | 0 |
| C | $w_{max}$ | 0 | $w_{max}$ | 0 |
| D | 0 | 0 | 0 | 0 |

For further evaluation, instead of the Gaussian noise term *e_i,h_*, we simulated another noise term 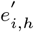 as noise derived from a gamma distribution (Fig. S2a). We simulated 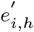 for each gene *h* by using a gamma distribution whose parameters were based on the seqFISH+ real dataset. This noise matrix was log-scaled and z-score normalized as 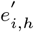.

In section 3.1, we present the application of CCPLS to this simulated dataset, including an evaluation of the results of cell type A. We first set the *w_max_* and *α* to 1 in Figure 2a and then changed in Figure S1a. Next, we changed the noise term *e_i,h_* with 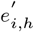 (Fig. S2). See text S2 in the Supplementary Information for the detailed procedures.

#### 2.3.2 SeqFISH+ real dataset

For the demonstration of CCPLS with a real dataset, we selected the seqFISH+ mouse somatosensory cortex dataset provided by Giotto (https://rubd.github.io/Giotto_site/articles/mouse_seqFISH_cortex_200914) (Dries *et al*., 2021b). This seqFISH+ dataset profiled 10,000 genes in 523 cells with 12 distinct cell types at a single-cell resolution by using a super-resolved imaging technique (Eng *et al*., 2019).

In section 3.2, we describe the results of applying CCPLS to this seqFISH+ real dataset. We further assigned the contributor cell types, which were the common neighboring cell types in each HVG cluster. We performed Gene Ontology (GO) enrichment analysis for each HVG cluster to investigate the associated biological insights. See text S2 in the Supplementary Information for the detailed procedures.

#### 2.3.3 Seq-Scope real dataset

For the demonstration of CCPLS with a real dataset, we selected the Seq-Scope colon dataset, which can be acquired from the repository of Deep Blue Data (https://doi.org/10.7302/cjfe-wa35) (Cho *et al*., 2021). This Seq-Scope dataset profiled 10,806 genes in 4,489 cells with nine distinct cell types at a single-cell resolution by using Illumina flow-cell-based spatial barcoding techniques (Cho *et al*., 2021).

In section 3.3, we describe the results of applying CCPLS to this Seq-Scope real dataset, including the assignment of contributor cell types and GO enrichment analysis according to the same procedure referred to in section 2.3.2.

## 3 Results

### 3.1 CCPLS yields an accurate estimation of cell-cell communications

To evaluate CCPLS performance, we prepared simulated data with parameters based on the seqFISH+ real dataset and then applied CCPLS to this simulated data as described in Section 2.3.1 (Fig. 2a). We obtained a heat map and bipartite graph, which indicate the coefficient as the direction and degree of regulation in the cell-cell communication for each combination of HVGs and neighboring cell types by CCPLS (Fig. 2 b-c). Cell types C and B exhibited upregulation of HVG clusters 1 and 2, respectively (Fig. 2 b-c), while cell types A and C cooperatively up-regulated HVG cluster 3 (Fig. 2b-c). Three cell types, A, B, and C, exhibited upregulation of HVG cluster 4, whose degree of cell-cell communications was relatively low (Fig. 2c).

We assigned the estimated HVG clusters 1-4 with the predefined clusters. The estimated HVG clusters 1, 2, and 3 corresponded to the predefined HVG clusters 3, 2, and 1, respectively. The estimated HVG cluster 4 was assigned as “others”. Note that CCPLS did not report HVGs whose filtered coefficients were all zero, which corresponded to the predefined HVG cluster 4 in this case. Based on the cluster assignment, we calculated ten indexes for performance evaluation (Fig. 2d). All the calculated indexes for CCPLS results were greater than 0.97, which indicated that CCPLS estimated the prepared coefficients and the predefined HVG clusters with high performance. We also compared CCPLS with the existing method Giotto findICG, which was outperformed by CCPLS in this simulation experiment (Dries *et al*., 2021b) (Fig. 2d). In HVG cluster 1 of the multiple-input case, the performance between the two methods was most different. Note that we did not compare CCPLS to the other methods, which do not estimate the effect of neighboring cell types (Arnol *et al*., 2019; Dries *et al*., 2021b; Hu *et al*., 2021a; Rao *et al*., 2021; Svensson *et al*., 2018; Tanevski *et al*., 2022; Velten *et al*., 2022; Zhu *et al*., 2021) (Table S1).

Next, we examined the performance limits of CCPLS by changing *w_max_* and *α*, the parameters for adjusting the degree of the cell-cell communications and Gaussian noise, respectively (Fig. S1a). According to the same procedures shown in Figure 2, we calculated the ten indexes (Fig. S1b). In each *w_max_*, these 10 indexes decreased when α increased. These ten indexes were greater than 0.97 for conditions 1-5 and 7, which indicated the estimations were successful. In conditions 6 and 8, these ten indexes were relatively lower and moderate. The estimation failed under only condition 3, for which the eight indexes were less than 0.02 (Fig. S1b).

We further investigated the difference between the failed and successful conditions by calculating the index of variance proportion (*VP*) as the ratio of the variance derived from the regression model term not but the Gaussian noise term. In each *w_max_*, *VP* decreased when *α* increased, which indicated that the higher α conditions were more severe, such that the variance associated with cell-cell communications was more difficult to distinguish from noise (Fig. S1c). The *VP* of the failed conditions were less than 0.04, while the *VP* of the successful conditions were greater than 0.61. The estimation did not work well under excessively severe *VP* conditions, while CCPLS estimated the prepared coefficients and the predefined HVG clusters with high performance in the relatively higher *VP* cases. Note that CCPLS did not estimate any HVGs and cell-cell communications from randomly generated data (data not shown).

We also generated another simulated dataset by replacing Gaussian noise *e_i,h_* with noise derived from gamma distribution 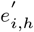 (See section 2.3.1) and evaluated the performance of CCPLS on the Gamma-noise simulated dataset. The result showed similar performance and tendency with the cases by using Gaussian noise (Fig. S2).

These simulation experiments showed that CCPLS yielded an accurate estimation of the MIMO system of cell-cell communications.

### 3.2 Application to seqFISH+ real dataset reveals biologically consistent insights associated with oligodendrocyte differentiation

To demonstrate CCPLS with a real dataset, we applied CCPLS to the seqFISH+ real dataset (Eng *et al*., 2019; Dries *et al*., 2021b). Procedures were as described in Section 2.3.2. Before we applied CCPLS, we examined the overlap of the extracted HVGs among all 12 of the cell types in the seqFISH+ real dataset (Fig. 3b). The mean value of the ratio of the overlapping HVGs was 0.25. The majority of the HVGs were different in each comparison, which indicated that the HVGs were uniquely characterized for each cell type.

**Fig. 3.**
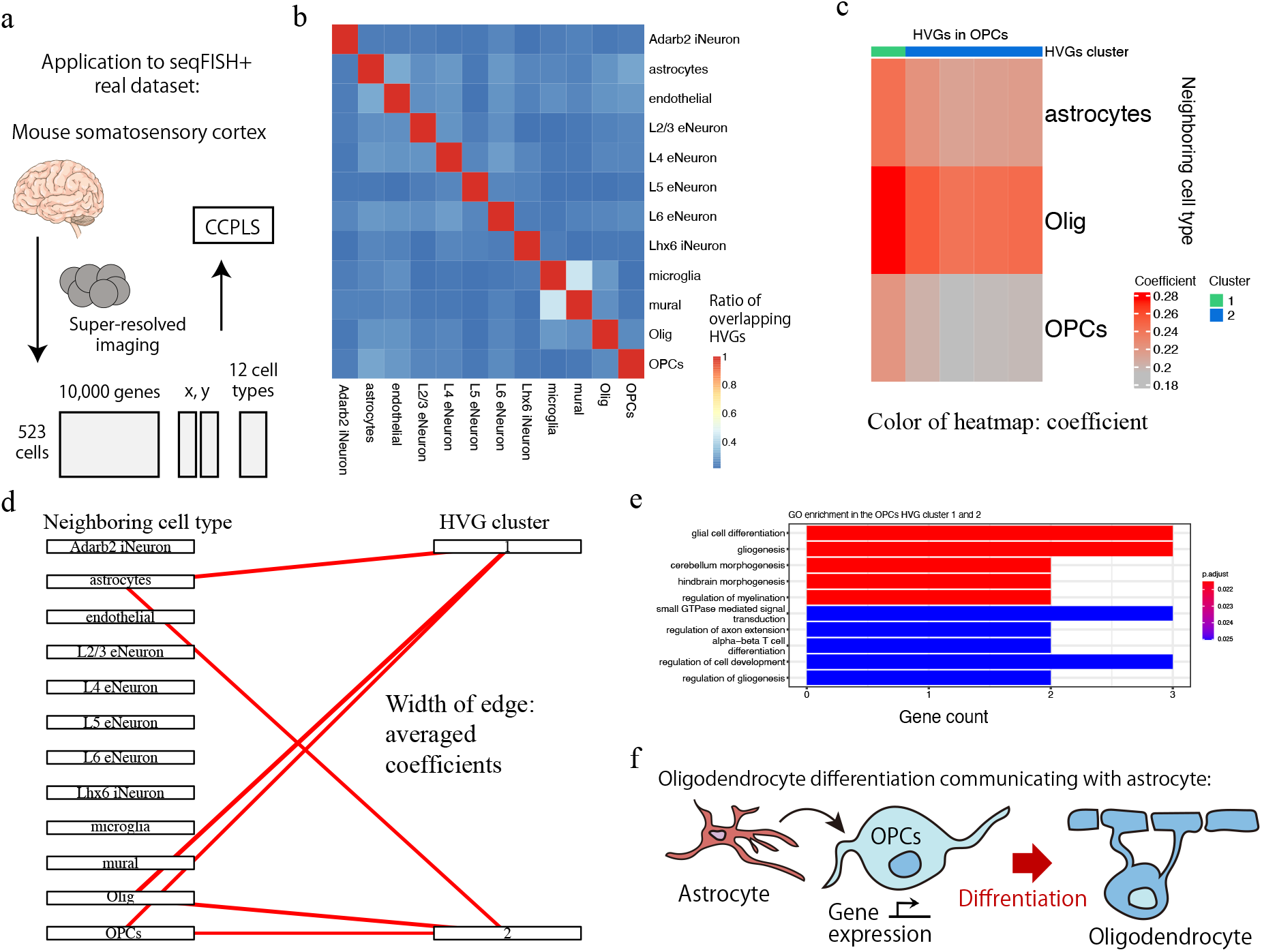
Application to the seqFISH+ real dataset. (a) Schematic illustration of the seqFISH+ dataset. (b) Overlap of highly variable genes (HVGs) between cell types. Rows and columns both correspond to cell types. The color of the heat map indicates the ratio of the overlapping HVGs. iNeuron: inhibitory neuron, endothelial: endothelial cells, eNeuron: excitatory neuron, mural: mural cells, Olig: oligodendrocyte. (c) Heat map of oligodendrocyte precursor cells (OPCs) generated by CCPLS. Rows correspond to neighboring cell types. Columns correspond to HVGs. The color of the heat map indicates the coefficient. (d) Bipartite graph of OPCs generated by CCPLS. The width of the edge indicates the averaged coefficients for each combination of HVG clusters and neighboring cell types. (e) Gene Ontology (GO) enrichment of HVG clusters 1 and 2 for OPCs. The top 10 GO terms are indicated. (f) Schematic illustration of the obtained biological insights on OPCs.

To estimate the cell-cell communications associated with these HVGs for each cell type, we applied CCPLS. We obtained the HVGs and the cellcell communications for six cell types out of all twelve cell types (Fig. 3c-d and S2-3). Focusing on the oligodendrocyte precursor cells (OPCs), two HVG clusters were estimated (Fig. 3c-d). We assigned the contributor cell types, which indicated that three cell types (astrocytes, oligodendrocyte (Olig), and OPCs) cooperatively up-regulated HVG clusters 1 and 2 in the OPCs.

In addition, we performed GO enrichment analysis to investigate associated biological insights. The GO enrichment analysis was performed on the mixed gene list of HVG clusters 1 and 2, as HVG clusters 1 and 2 have the same contributor cell types. The GO enrichment of HVG clusters 1 and 2 in OPCs indicated the 25 enriched GO terms (Fig. S5). The top 10 GO terms are indicated in Figure 4e (Fig. 3e). GO terms related to differentiation were enriched, such as “glial cell differentiation,” indicating that differentiation of OPCs was induced by communicating with astrocytes, Olig, and OPCs itself (Fig. 3f). Additionally, we plotted the positions of the cells in the 2D space and showed the expression level of *Mag* gene, which is a genes annotated with the GO term “glial cell differentiation” (Fig. S3). This visualization showed that the OPCs near astrocytes, Olig, and OPCs showed high expression of *Mag* genes.

**Fig. 4.**
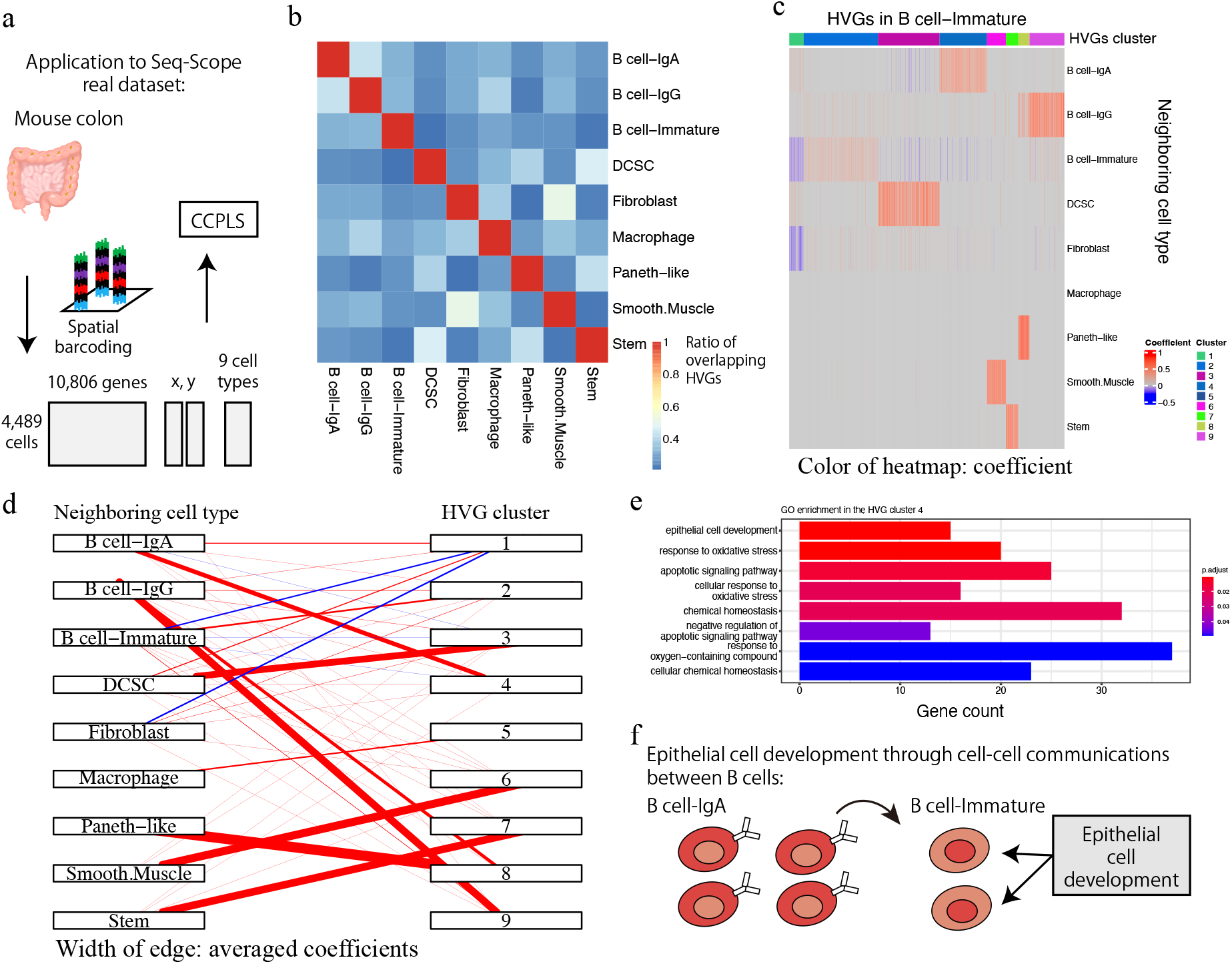
Application to the Seq-Scope real dataset. (a) Schematic illustration of the Seq-Scope dataset. (b) Overlap of highly variable genes (HVGs) between cell types. Rows and columns both correspond to cell types. The color of the heat map indicates the ratio of the overlapping HVGs. (c) Heat map of the immature B cell generated by CCPLS. Rows correspond to neighboring cell types. Columns correspond to HVGs. The color of the heat map indicates the coefficient. (d) Bipartite graph of the immature B cell generated by CCPLS. The width of each edge indicates the average coefficients for each combination of HVG clusters and neighboring cell types. (e) Gene Ontology (GO) enrichment of HVG cluster 4 of the immature B cell. (f) Schematic illustration of the obtained biological insights on the immature B cell.

These obtained insights were principally consistent with previous *in vivo* mice experiments showing that astrocytes induce the differentiation of OPCs into oligodendrocytes (Nutma *et al*., 2020) (Fig. 3f). In contrast, when we applied Giotto findICG (Dries *et al*., 2021b) on the SeqFISH+dataset, few overlaps were observed between the detected genes by Giotto findICG and CCPLS (Fig. S4). Moreover, for the cell type pairs where the receiver cell type is OPCs and the neighboring cell type is astrocyte, Olig, or OPCs itself, we found no GO enrichment in the genes detected by Giotto findICG (data not shown). Furthermore, these genes showed fewer overlap with the genes annotated with the GO term “glial cell differentiation” than CCPLS (Fig. S5). Considering that CCPLS can handle combinations of neighboring cell types, these results suggest that CCPLS detects cell-cell communications that cannot be detected by Giotto findICG.

The heat map, bipartite graph, contributor cell type assignment, and GO enrichment analysis results for all cell types are shown in the Supplementary Information (Fig. S6-9, respectively). This application to the seqFISH+ real dataset indicated the capability of CCPLS to extract biologically interpretable insights.

### 3.3 Application to Seq-Scope real dataset suggests epithelial cell development in immature B cell

For the further demonstration of CCPLS using a real dataset, we applied CCPLS to the Seq-Scope real dataset (Cho *et al*., 2021). The procedures followed are as described in Section 2.3.3. Before we applied CCPLS, we examined the overlapping of the extracted HVGs between all nine cell types in the Seq-Scope real dataset (Fig. 4b). The mean value of the ratio of the overlap HVG was 0.31. The majority of the HVGs differed between each comparison, which indicated that the HVGs were uniquely characterized for each cell type.

To estimate cell-cell communications associated with these HVGs for each cell type, we applied CCPLS. We obtained the HVGs and the cell-cell communications for eight cell types out of all nine cell types (Fig. 4c-d and S6-7). Focusing on the immature B cell, nine HVG clusters were estimated (Fig. 4c-d). We assigned the contributor cell types, which indicated that the immature B cell and the Fibroblast cell types cooperatively down-regulated HVG cluster 1 in the immature B cell. In HVG clusters 2 to 9, there was one contributing cell type associated with up-regulation. The contributor cell types in clusters 2 to 9 were immature B cell, DCSC, IgA B cell, Macrophage, Smooth Muscle, Stem, Paneth-like, and IgG B cell, respectively.

In addition, we performed GO enrichment analysis to investigate potential biological insights. The GO enrichment of cluster 4 for the immature B cell indicated, among the eight enriched GO terms, “epithelial cell development” was the most enriched term (Fig. 4e). We further plotted the positions of the cells in the 2D space and visualized the expression level of *Gpx1* gene, a gene annotated with the GO “epithelial cell development”, in B cell-Immature (Fig. S10), showing that the B cell-Immature cells on the boundary between B cell-Immature and B cell-IgA showed high expression of *Gpx1* genes.

These results suggested that epithelial cell development of the immature B cell occurred through communicating with IgA B cells (Fig. 4f), which have not been reported previously to the best of our knowledge (Goto, 2019) and would be deserving of further investigation. In contrast, when we applied Giotto findICG (Dries *et al*., 2021b) on the Seq-Scope dataset, less than 30 percent of genes were overlapped on average (Fig. S11). For the cell type pairs where the receiver cell type is immature B cell and the neighboring cell type is IgA B cell, we found no GO enrichment in genes detected by Giotto findICG (data not shown). Furthermore, we examined the number of overlap between genes of GO “epithelial cell development”, genes detected by CCPLS, and genes detected by Giotto findICG in those cell type pairs (Fig. S12), revealing the genes annotated wtih the GO term “epithelial cell development” were detected by only CCPLS (15 of 15 genes).

The heat map, bipartite graph, contributor cell type assignment, and GO enrichment analysis results for all the cell types are shown in the Supplementary Information, respectively (Fig. S13-16). The application to the Seq-Scope real dataset demonstrate the capability of CCPLS to extract biologically interpretable insights.

### 4 Discussion

In this study, we propose CCPLS (Cell-Cell communications analysis by Partial Least Square regression modeling), which is a statistical framework for identifying the MIMO system of cell-cell communications, i.e., gene expression regulation by multiple neighboring cell types, based on the spatial transcriptome data at a single-cell resolution. CCPLS estimates regulation on cell-to-cell expression variation of HVGs via cell-cell communications not limited to ligand-receptor pairs. The core of CCPLS is PLS regression modeling performed by setting the HVGs expression values as the responsive variables and the neighboring cell-type scores as the explanatory variables for each cell type. After the statistical evaluation, CCPLS performs HVG clustering, which is reported with coefficients as the direction and degree of regulation in the cell-cell communication. Evaluation using simulated data showed accurate estimation by CCPLS. The application of CCPLS to the two real spatial transcriptome datasets demonstrated the capability to extract biologically interpretable insights.

CCPLS differs from the existing method Giotto findICG in that CCPLS quantitatively evaluates cell-cell communications as a MIMO system by including cooperation of neighboring cell types, which may have revealed the biological insights derived from the applications to the two real datasets. In the simulation experiments, CCPLS outperformed Giotto findICG, especially with regard to HVG cluster 1, which is a multiple-input case.

The application of CCPLS to the seqFISH+ real dataset showed that the differentiation in the OPCs occurred through communicating with astrocytes, Olig, and OPCs itself, which was principally consistent with previous *in vivo* mice experiments and not identified by Giotto findICG (Dries *et al*., 2021b; Nutma *et al*., 2020). The application to the Seq-Scope real dataset suggested that epithelial cell development is induced in the immature B cell by communicating with the IgG B cells, which has not been reported previously to the best of our knowledge (Goto, 2019). Accordingly, further experimental follow-up studies are merited. Other existing methods, which do not focus on the effect of neighboring cell types, may have missed these findings (Arnol *et al*., 2019; Dries *et al*., 2021b; Hu *et al*., 2021a; Svensson *et al*., 2018; Tanevski *et al*., 2022; Velten *et al*., 2022; Zhu *et al*., 2021). As another characteristic, CCPLS focuses on HVGs not limited to ligand-receptor pairs, which may have also enabled it to reveal the biological insights noted above.

The exploration of cell-cell communications in combination with spatial transcriptome data is still largely incomplete. CCPLS would be effective for exploring drug targets as in the case of the PD1 pathway, which led to a breakthrough drug discovery (Sharpe and Pauken, 2018). For example, the treatment of cold tumors, which are composed of tumor cells under a severe microenvironment with few neighboring T cells, is still considered to be a particular challenge (Haanen, 2017). The severe tumor progression possibly derived from the cold tumor-specific combinations of HVGs and neighboring cell types, which thus may be candidates for drug targeting. Specifically, CCPLS would be suitable in cases where cell arrangement causes gene expression changes.

The limitations and perspectives in this study are as follows. The first is the model assumption of CCPLS. In this study, we assumed that highly variable genes can be explained by a linear combination of neighboring cell-type scores. Several studies exemplify changes in gene expression is associated with or induced by the placement of surrounding cell types in multicellular systems (Colombo and Cattaneo, 2021; Haanen, 2017; Hui and Bhatia, 2007). Such a method of detecting the effect of surrounding cell types on gene expression could be biologically useful. For CCPLS, we decided to assume a linear combination of neighboring cell-type scores is a simple model, which is easy to interpret and computationally tractable. However, it is possible that the refinement of the mathematical model of the effect of cell type on gene expression (e.g., nonlinear relationships among cell types) is an important issue to be addressed in the future.

The second is the potential modifications of CCPLS. Although CCPLS employed PLS regression, the other sparse regression can be candidates such as lasso or ridge regression, whose implementations and evaluations are an intended focus of our future work (Tibshirani, 1996). CCPLS employed a Gaussian kernel for calculating the neighboring cell-type score similar to the previous studies (Arnol *et al*., 2019; Dang *et al*., 2020). However, the other kernels such as Matern kernel or exponential kernel can be possible, whose implementations and evaluations are also an intended focus of our future work (Pustokhina *et al*., 2021). CCPLS selected *dist*0 as the minimum value of the Euclidean distance of all the combinations of two cells, while the other selection procedures may work. The selection of parameter *dist*0 is also one of our future works.

The third is the incorporation of cell-cell communications other than the neighboring cell effects. CCPLS explicitly handles combinations of neighboring cell types as advantages compared to the existing methods (Arnol *et al*., 2019; Dries *et al*., 2021b; Hu *et al*., 2021a; Rao *et al*., 2021; Svensson *et al*., 2018; Tanevski *et al*., 2022; Velten *et al*., 2022; Zhu *et al*., 2021) (Table S1), while CCPLS neither extracts nonlinear relationships nor estimates the cell-cell communications derived from factors beyond the neighboring cell-type score, such as ligand-receptor pairs and cellcell communications via long-range effect (Dries *et al*., 2021a; Fechner and Goodman, 2018; Longo *et al*., 2021; Palla *et al*., 2022; Rao *et al*., 2021; Sapir and Tzlil, 2017). Handling molecular pathways involved in the MIMO system is an intended focus of our future work.

The fourth is the extension of CCPLS to other types of spatially resolved omics data. CCPLS does not receive as input spatial transcriptome data with a multiple-cell resolution as is characteristic of 10x Genomics Visium and Slide-seq datasets (Rodriques *et al*., 2019; Stickels *et al*., 2021; Asp *et al*., 2020). Expansions of CCPLS to address these limitations are an intended focus of our future work. In addition, the responsive expression values can be replaced by the other omics data such as spatial metabolome or proteome data, which is also an intended focus of our future work (Alexandrov, 2020; Bhatia *et al*., 2021; Dyring-Andersen *et al*., 2020; Geier *et al*., 2020; Liu *et al*., 2020; Seydel, 2021).

Currently, collaborative consortiums are accumulating large-scale spatial transcriptome data (BRAIN Initiative Cell Census Network (BICCN), 2021; Zhang *et al*., 2021; Regev *et al*., 2017), and CCPLS can be readily applied to these large-scale datasets. CCPLS is provided as a package in the R statistical computing environment, and its calculation time was less than 10 minutes for each demonstration presented in this study. In summary, CCPLS provides novel insights into regulation on cell-to-cell expression variability of HVGs via cell-cell communications.

## Acknowledgements

We thank Dr. Wataru Iwasaki, Dr. Hirotaka Matsumoto, Dr. Tsukasa Fukunaga and Dr. Kentaro Kawata for critical reading of this manuscript. The computational analysis of this work was performed in part with the support of the supercomputer system of the National Institute of Genetics (NIG), Research Organization of Information and Systems (ROIS). We thank kango-roo (www.kango-roo.com) for providing illustrations.

## Software and data availability

The R package is available at https://github.com/bioinfo-tsukuba/CCPLS. The data are available at https://github.com/bioinfo-tsukuba/CCPLS_paper.

