## Supplementary Information for "CCPLS reveals cell-type-specific spatial dependence of transcriptomes in single cells"

Contents:

Text S1. Filtering of coefficients in CCPLS

Text S2. Additional descriptions of datasets

Table S1. Computational methods for spatial transcriptome data

Figure S1. Evaluation using the simulated datasets across parameters

Figure S2. Evaluation using the noise derived from gamma distribution.

Figure S3. Spatial distribution of Mag expression in Oligodendrocyte Precursor cells (OPCs).

Figure S4. Comparison between Giotto findICG and CCPLS in the seqFISH+ real dataset.

Figure S5. Number of overlaps between genes of GO "glial cell differentiation", genes detected by CCPLS, and genes detected by Giotto findICG.

Figure S6. Heatmap of all the cell types in the seqFISH+ real dataset

Figure S7. Bipartite graph of all the cell types in the seqFISH+ real dataset

Figure S8. Classification of the contributor cell types in the seqFISH+ real dataset

Figure S9. GO enrichment of all the cell types in the seqFISH+ real dataset

Figure S10. Spatial distribution of Gpx1 expression in B cell-Immature.

Figure S11. Comparison between Giotto findICG and CCPLS in the Seq-Scope read dataset.

Figure S12. Number of overlaps between genes of GO "epithelial cell development", genes detected by CCPLS, and genes detected by Giotto findICG.

Figure S13. Heatmap of all the cell types in the Seq-Scope real dataset

Figure S14. Bipartite graph of all the cell types in the Seq-Scope real dataset

Figure S15. Classification of the contributor cell types in the Seq-Scope real dataset

Figure S16. GO enrichment of all the cell types in the Seq-Scope real dataset

### Text S1. Filtering of coefficients in CCPLS

In filtering step (i), CCPLS calculates *p*-values for $w_{f,h,c}^{(m)}$ of each component $c$, which is an element of $w_{f,h}^{(m)}$ and is obtained by PLS regression modeling. The *p*-values are calculated by *t*-tests of factor loadings (Yamamoto *et al.*, 2014), which correspond to the *p*-values of the Pearson correlation coefficient calculated by

$$\mathrm{corr}\left(\mathbf{x}_\mathrm{f}^{(\mathrm{m})}, \mathbf{t}_\mathrm{c}^{(\mathrm{m})}\right) = \frac{\mathbf{x}_\mathrm{f}^{(\mathrm{m})}\mathbf{t}_\mathrm{c}^{(\mathrm{m})}/(\mathrm{N}^{(\mathrm{m})}-1)}{\sqrt{\mathrm{var}\left(\mathbf{x}_\mathrm{f}^{(\mathrm{m})}\right)}\sqrt{\mathrm{var}\left(\mathbf{t}_\mathrm{c}^{(\mathrm{m})}\right)}} \ ,$$

$$\mathrm{corr}\left(\mathbf{y}_\mathrm{h}^{(\mathrm{m})}, \mathbf{u}_\mathrm{c}^{(\mathrm{m})}\right) = \frac{\mathbf{y}_\mathrm{h}^{(\mathrm{m})}\mathbf{u}_\mathrm{c}^{(\mathrm{m})}/(\mathrm{N}^{(\mathrm{m})}-1)}{\sqrt{\mathrm{var}\left(\mathbf{y}_\mathrm{h}^{(\mathrm{m})}\right)}\sqrt{\mathrm{var}\left(\boldsymbol{u}_\mathrm{c}^{(\mathrm{m})}\right)}} \ ,$$

where $\mathrm{corr}()$ and $\mathrm{var}()$ denote calculations of a Pearson correlation coefficient and its variance, respectively. The vectors $\mathbf{x}_\mathrm{f}^{(\mathrm{m})}$ and $\mathbf{y}_\mathrm{h}^{(\mathrm{m})}$ are preprocessed scores of neighboring cell type $f$ and preprocessed expression values of HVG $h$ of cells $i$ within cell type $m$. The vectors $\mathbf{t}_\mathrm{c}^{(\mathrm{m})}$ and $\mathbf{u}_\mathrm{c}^{(\mathrm{m})}$ are scores of $c$-th component obtained by PLS regression modeling. These *p*-values are false discovery rate (FDR)-adjusted as $q_\mathrm{f,c}^{(\mathrm{m})}$ and $q_\mathrm{h,c}^{(\mathrm{m})}$ by the Benjamini–Hochberg (BH) method, respectively (Benjamini and Hochberg, 1995). CCPLS filters out the coefficient $\mathrm{w}_\mathrm{f,h,c}^{(\mathrm{m})}$ whose adjusted *p*-values $q_{f,c}^{(m)}$ or $q_{h,c}^{(m)}$ are greater than or equal to $\alpha$ and then returns the coefficient $\mathrm{w'}_\mathrm{f,h}^{(\mathrm{m})} = \Sigma_\mathrm{c} \mathrm{w}_\mathrm{f,h,c}^{\prime(\mathrm{m})}$ as follows:

$$\begin{cases} w_{f,h,c}^{\prime(m)} = w_{f,h,c}^{(m)} & \text{if } q_{f,c}^{(m)} < \alpha \vee q_{h,c}^{(m)} < \alpha \\ w_{f,h,c}^{\prime(m)} = 0 & otherwise \end{cases} .$$

In this study, we set $\alpha$ to 0.05.

In filtering step (ii), CCPLS filters out the statistically non-significant coefficient $\mathrm{w}_\mathrm{f}^{\prime(\mathrm{m})}$ by using a non-parametric test. CCPLS uses the coefficients $\mathrm{w}_\mathrm{f,h}^{(\mathrm{m})}$ which are statistically non-significant genes in all the neighboring cell types $f$ in the step (i) as a null distribution. For each neighboring cell type $f$, CCPLS calculates the adjusted *p*-values $q_\mathrm{f,h}^{\prime(\mathrm{m})}$ by the BH method (Benjamini and Hochberg, 1995). For each neighboring cell type $f$, CCPLS filters out the coefficient $\mathrm{w}_\mathrm{f,h}^{\prime(\mathrm{m})}$ whose adjusted *p*-values $q_\mathrm{f,h}^{\prime(\mathrm{m})}$ are greater than or equal to $\alpha$ and then returns $\mathrm{w}_\mathrm{f,h}^{\prime\prime(\mathrm{m})}$ as follows:

$$\begin{cases} w_{f,\mathrm{h}}^{\prime\prime(m)} = w_{f,\mathrm{h}}^{\prime(m)} & if\ q_{f,h}^{\prime(m)} < \alpha \\ w_{f,\mathrm{h}}^{\prime\prime(m)} = 0 & otherwise \end{cases} .$$

### Text S2. Additional descriptions of datasets

#### S2.1 Simulated dataset

We prepared the cell type label vector $\boldsymbol{L}$ by substituting cell types A-D into the cell type label vector of the seqFISH+ real dataset as follows:

- A: L5 eNeuron
- B: L6 eNeuron
- C: Olig
- D: The other nine cell types

In section 3.1, we assigned the correspondence between the estimated and predefined highly variable gene (HVG) clusters based on whether greater than half of genes had estimated coefficients corresponding to each flag in the following table, respectively. We assigned clusters to the group "others" if they were not assigned to any of the predefined clusters.

| Neighboring cell type | Flag of predefined cluster 1 | Flag of predefined cluster 2 | Flag of predefined cluster 3 | Flag of predefined cluster 4 |
|---|---|---|---|---|
| A | Significant | Not significant | Not significant | Not significant |
| B | Not significant | Significant | Not significant | Not significant |
| C | Significant | Not significant | Significant | Not significant |
| D | Not significant | Not significant | Not significant | Not significant |

For Giotto findICG, we assigned HVG cluster 1 if the sender cell type was "A" or "C," and HVG clusters 2-4 if the sender cell type was "B," "C," or none-detected, respectively.

For CCPLS and Giotto findICG, based on these assignments, we calculated the adjusted Rand index, precision, and recall for each HVG cluster. We calculated Pearson correlation coefficients from estimated coefficients not divided according to each assignment. We also calculated the index of variance proportion ($VP$) as follows:

$$\mathrm{VP} = \frac{1}{\mathrm{H}} \Sigma_{\mathrm{h}} \{ \frac{\mathrm{var}\left(\mathrm{x}_{\mathrm{i,f}} \mathrm{w}_{\mathrm{f,h}}^{(\mathrm{A})}\right)}{\mathrm{var}\left(\mathrm{x}_{\mathrm{i,f}} \mathrm{w}_{\mathrm{f,h}}^{(\mathrm{A})} + \alpha \mathrm{e}_{\mathrm{i,h}}\right)} \} \; .$$

#### S2.2 SeqFISH+ real dataset

In section 3.2, we assigned the contributor cell types, which were the common neighboring cell types for each HVG cluster. If the coefficient vector $\boldsymbol{w}_{\mathrm{f}}^{(\mathrm{m})}$ was significant in more than half of genes relative to the neighboring cell type $f$, we assigned it as the contributor cell type.

We performed Gene Ontology (GO) enrichment analysis of biological processes for each HVG cluster. If the distinct HVG clusters within the same cell type had the same contributor cell types, the genes were merged. We performed a hypergeometric test and extracted the significant GO terms with adjusted $p$- values less than 0.05 based on the BH method (Benjamini and Hochberg, 1995). We selected background genes whose raw expression values were greater than 0 within each cell type.

### Table S1. Computational methods for spatial transcriptome data

| Method | Reference | Effect of neighboring cell types | MIMO system | Descriptions |
|---|---|---|---|---|
| CCPLS | This study | Consider | Consider | Estimation of regulation on highly variable genes by multiple neighboring cell types based on PLS regression modeling. |
| Giotto findICG | Dires et al., 2021b | Consider | Not consider | Estimation of genes influenced by neighboring cell type based on spatial permutation test. |
| Giotto spatCellCellcom | Dires et al., 2021b | Consider | Not consider | Estimation of genes influenced by neighboring cell type based on permutation test. |
| SptialDE | Svensson et al., 2018 | Not consider | Not consider | Estimation of spatially variable genes based on Gaussian process regression modeling of spatial gene expression. |
| TrendSceek | Edsgärd et al., 2018 | Not consider | Not consider | Estimation of spatially variable genes based on marked point process modeling of spatial gene expression. |
| SPARK-X | Zhu et al., 2021 | Not consider | Not consider | Estimation of spatially variable genes based on spatial kernels and non-parametric modeling of spatial gene expression. |
| Giotto BinSpect | Dries et al., 2021b | Not consider | Not consider | Estimation of spatially variable genes based on enrichment analysis of spatially high expression cells after binarization. |
| Giotto SilhouetteRank | Dries et al., 2021b | Not consider | Not consider | Estimation of spatially variable genes by silhouette score per gene based on spatial distribution of two cells. |
| SVCA | Arnol et al., 2019 | Not consider | Not consider | Estimation of spatial variance sources of individual gene based on Gaussian process regression modeling of spatial gene expression. |
| SpaGCN | Hu et al., 2021a | Not consider | Not consider | Estimation of spatial domain and genes expression patterns based on graph convolutional network analysis. |
| MEFISTO | Velten et al., 2022 | Not consider | Not consider | Estimation of spatial gene expression patterns based on factor analysis. |
| MISTy | Tanevski et al., 2022 | Not consider | Not consider | Estimation of gene-gene relationships from different spatial views: intrinsic, local niche view, the broader, tissue view, or others. |

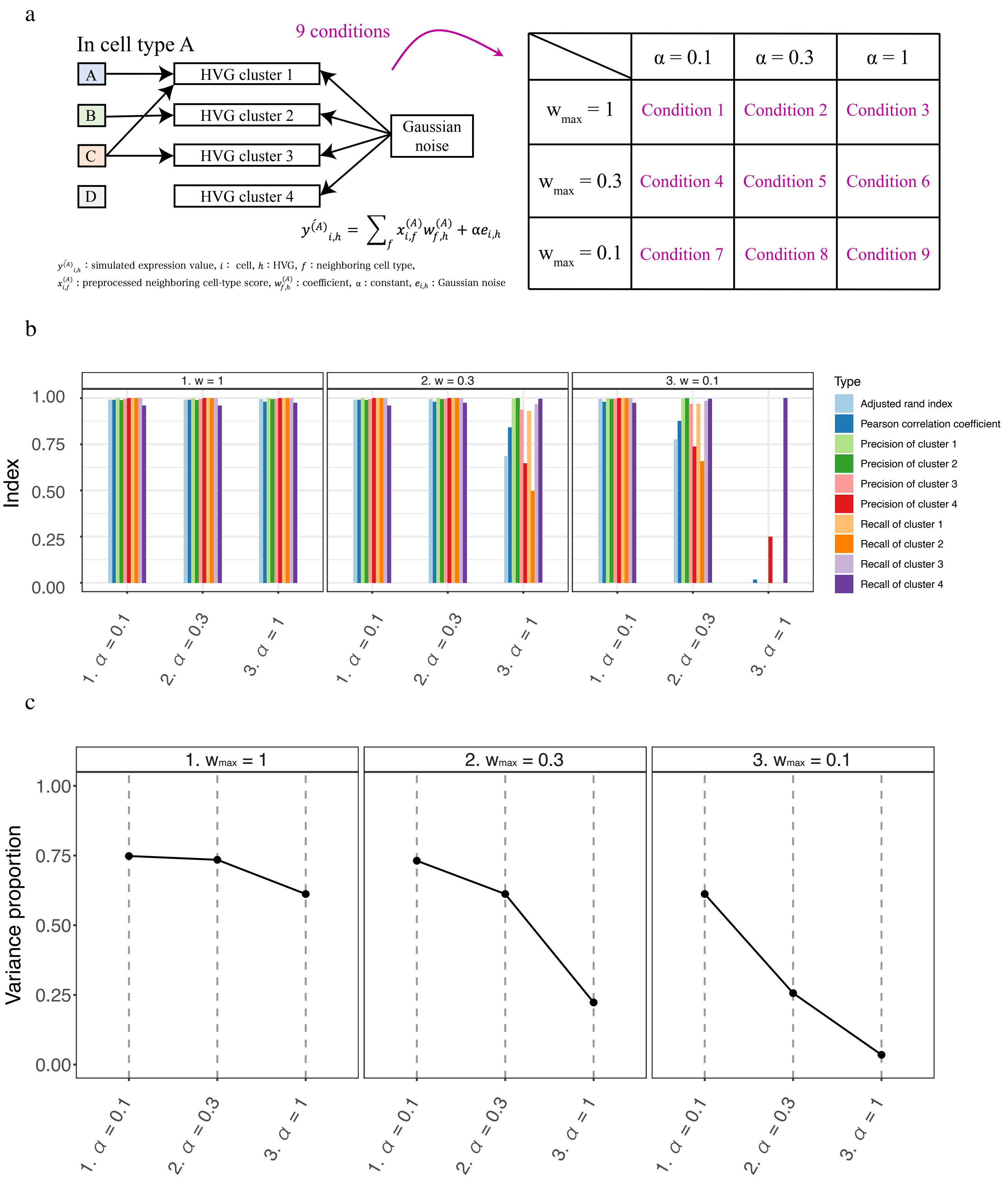


Figure S1. Evaluation using the simulated datasets across parameters. (a) Schematic illustration of the simulation settings with the changed parameters. Note that condition 3 corresponds to the condition in Figure 2. (b) Performance indexes in each condition. The value of each index is indicated along the y-axis, while each condition is arranged along the x-axis grouped within each $w_{max}$ value. The color indicates the index type. (c) Variance proportion in each condition. The variance proportion is indicated along the y-axis, while each condition is arranged along the x-axis grouped within each $w_{max}$ value.

a

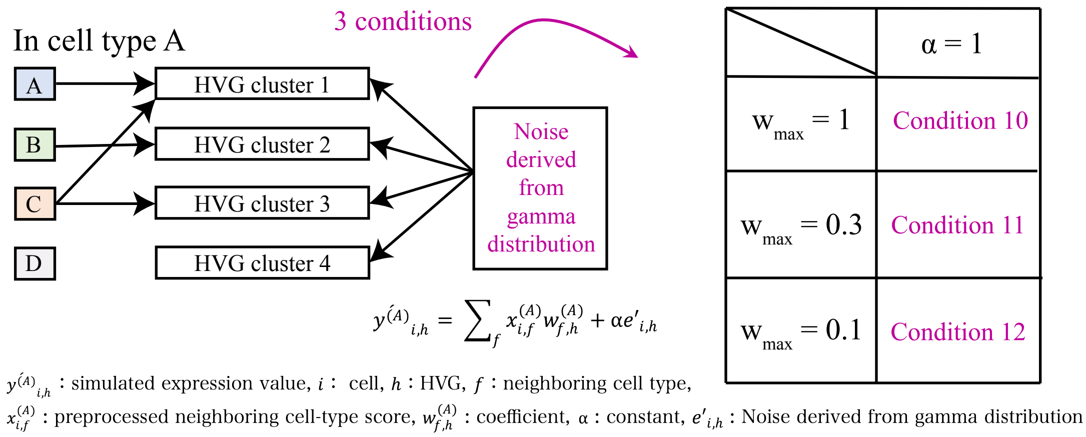


b

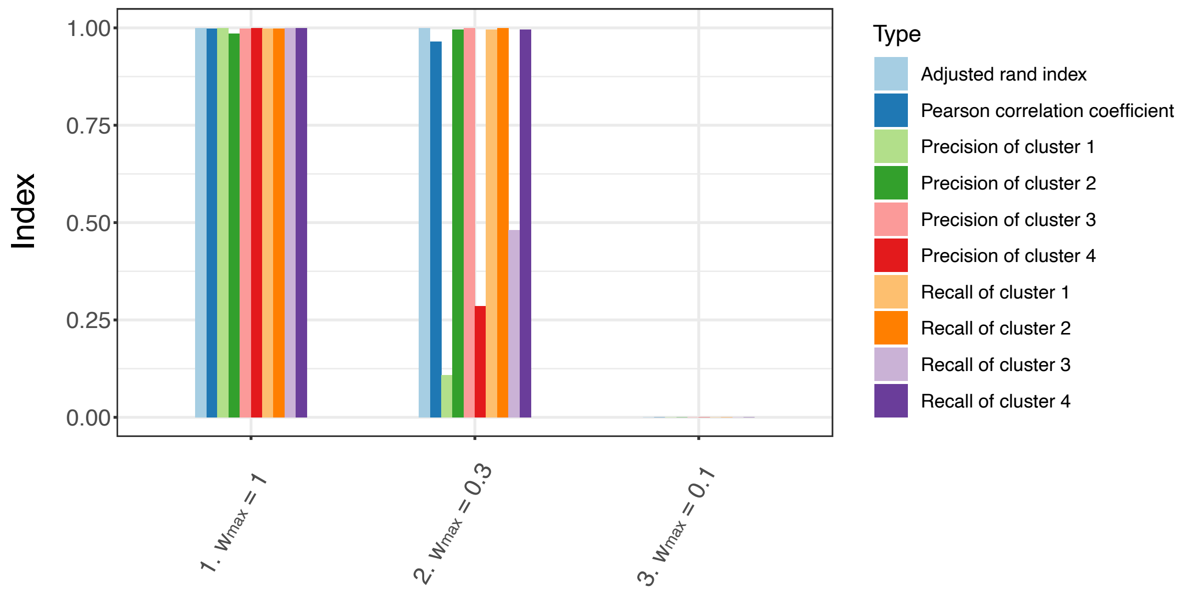


c

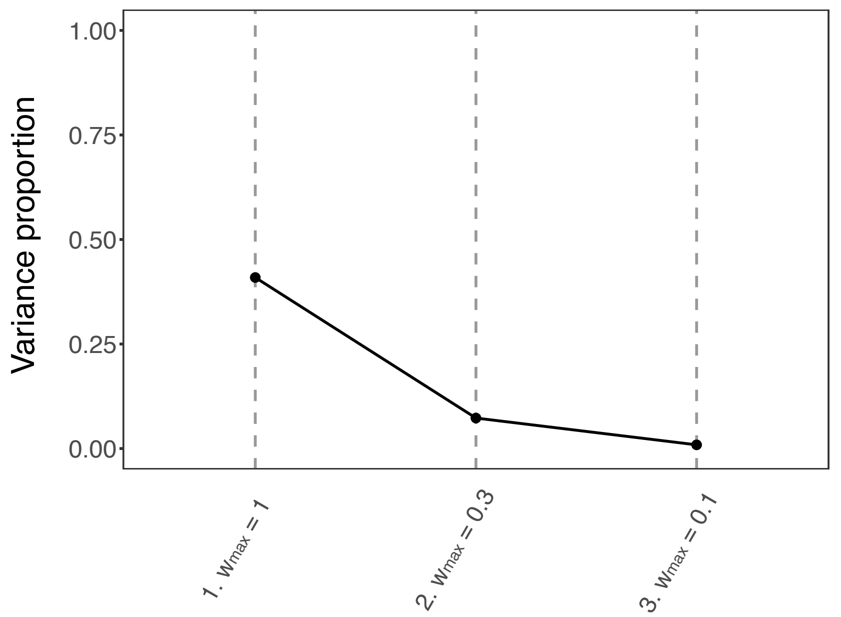


Figure S2. Evaluation using the noise derived from gamma distribution. (a) Schematic illustration of the simulation settings with the changed parameters. Note that we only replaced the Gaussian noise with the noise derived from gamma distribution compared with the Figure 2 and Figure S1. (b) Performance indexes in each condition. The value of each index is indicated along the y-axis, while each condition is arranged along the x-axis. The color indicates the index type. (c) Variance proportion in each condition. The variance proportion is indicated along the y-axis, while each condition is arranged along the x-axis grouped within each $w_{max}$ value.

### Apllication to the seqFISH+ real dataset:

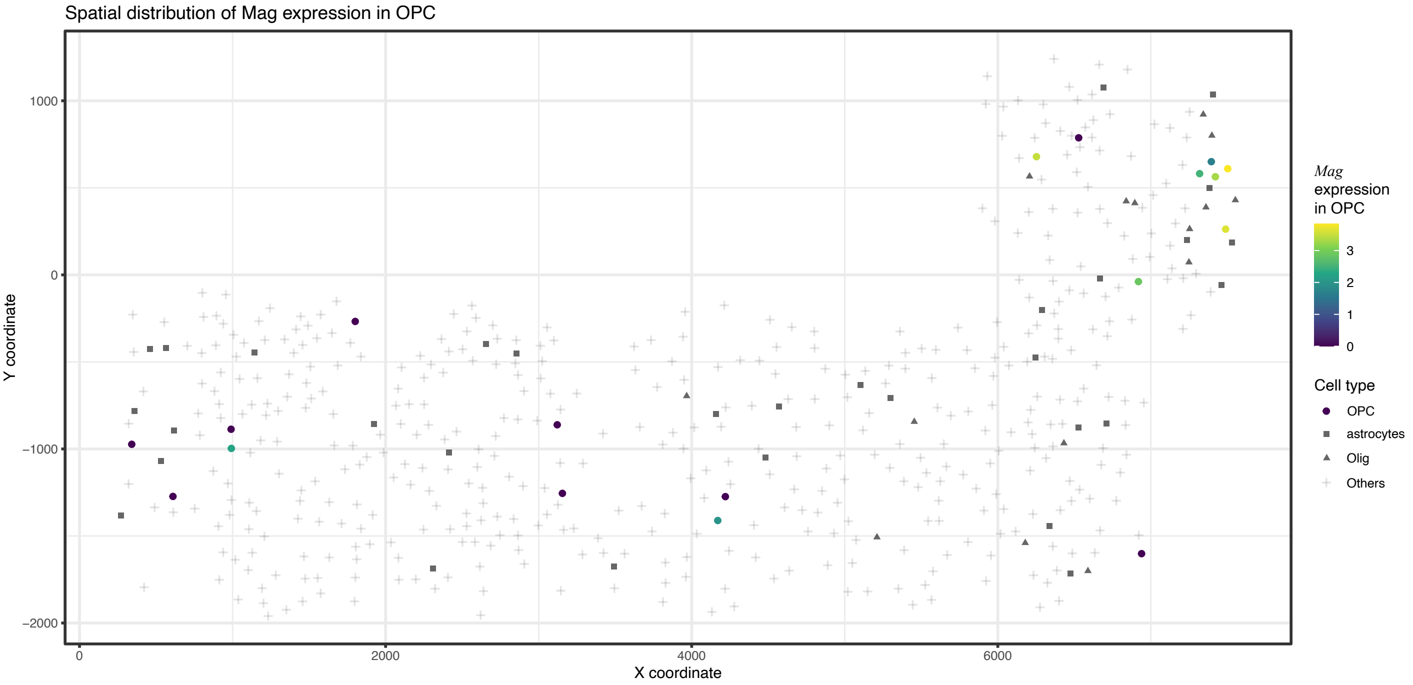


Figure S3. Spatial distribution of *Mag* expression in Oligodendrocyte Precursor cells (OPCs). The shapes indicate cell types. The color in the circles indicates values of *Mag* expression.

### Application to the seqFISH+ real dataset:

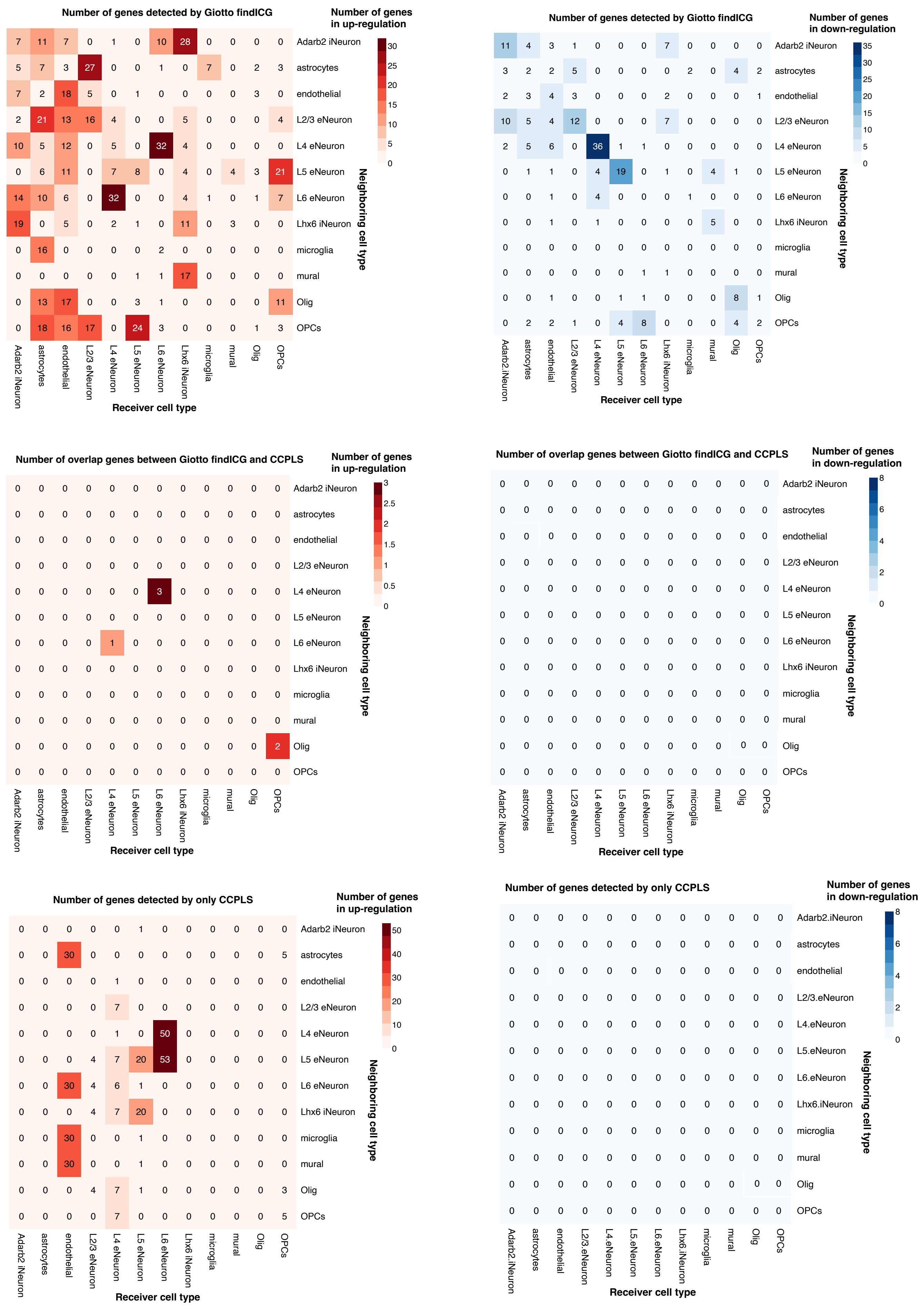


Figure S4. Comparison between Giotto findICG and CCPLS in the seqFISH+ real dataset. Note that no down-regulated genes were found for CCPLS (Fig. 3c and Fig. S6).

<u>Apllication to the seqFISH+ real dataset:</u>

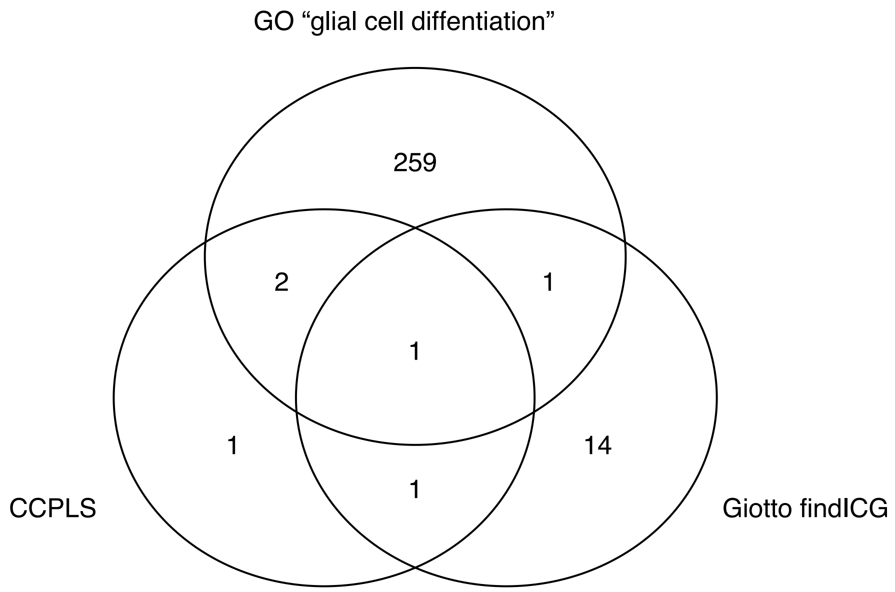


Figure S5. Number of overlaps between genes of GO “glial cell differentiation”, genes detected by CCPLS, and genes detected by Giotto findICG. Note that we extracted genes in Oligodendrocytes Precursor Cells (OPCs) up-regulated by astrocytes, Oligodendrocytes (Olig), or OPCs as to CCPLS and Giotto findICG in this venn diagram.

#### Apllication to the seqFISH+ real dataset:

Color of heatmap: coefficient

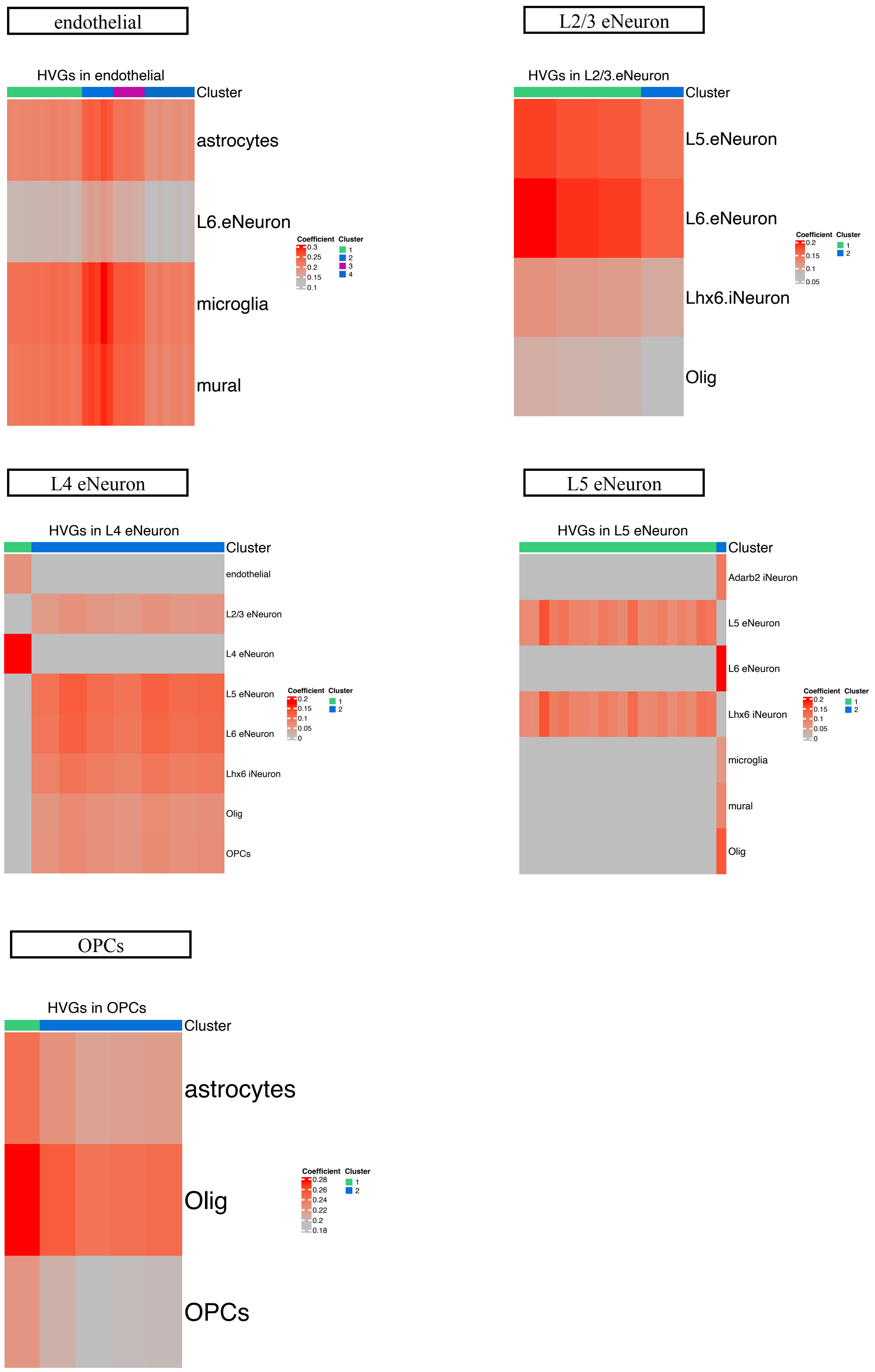


Figure S6. Heat map generated by CCPLS of all the cell types in the seqFISH+ real dataset. Rows and columns correspond to neighboring cell types and highly variable genes (HVGs), respectively. The color of the heat map indicates the coefficient. The heatmap of oligodendrocyte precursor cells (OPCs) is the same as that shown in Figure 3c.

### Application to the seqFISH+ real dataset:

Width of edge: averaged coefficients

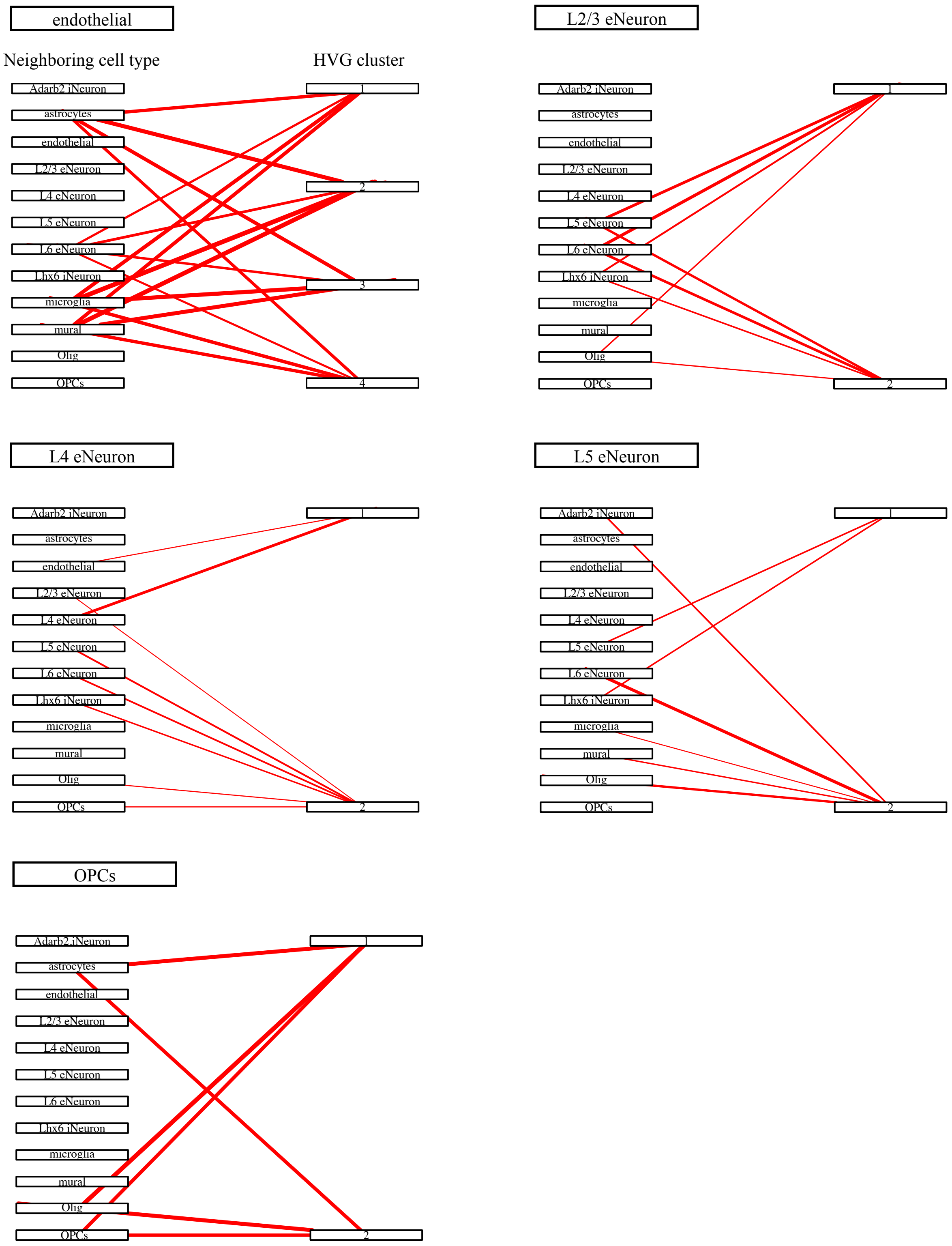


Figure S7. Bipartite graph generated by CCPLS of all the cell types in the seqFISH+ real dataset. The width of each edge indicates the averaged coefficients in each combination of highly variable gene (HVG) clusters and neighboring cell types. The bipartite graph of oligodendrocyte precursor cells (OPCs) is as the same as that shown in Figure 3d.

Application to the seqFISH+ real dataset:

Red: contributor cell type   Blue: non-contibutor cell type  Row: neighboring cell type   Column: HVG cluster

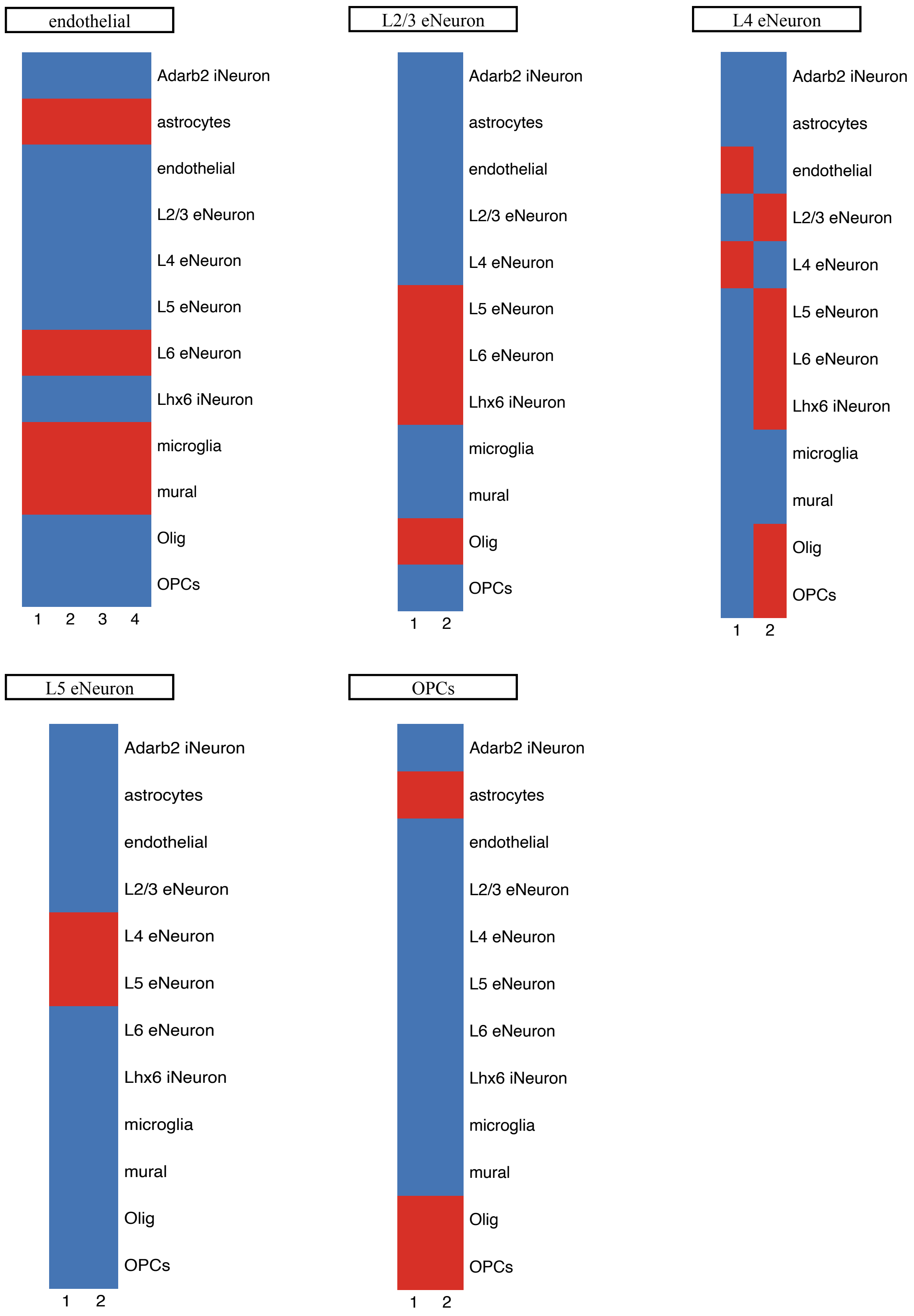


Figure S8. Contributor cell type in the seqFISH+ dataset. The color of the heat map corresponds to the binary value indicating whether the neighboring cell type is a contributor cell type or not. Red and blue indicate the contributor and non-contributor cell types, respectively. Rows and columns correspond to cell types and highly variable gene (HVG) clusters, respectively.

### Application to the seqFISH+ real dataset:

Row: GO term   Column: gene count
Color of bar graph: adjusted *p*-value

L2/3 eNeuron

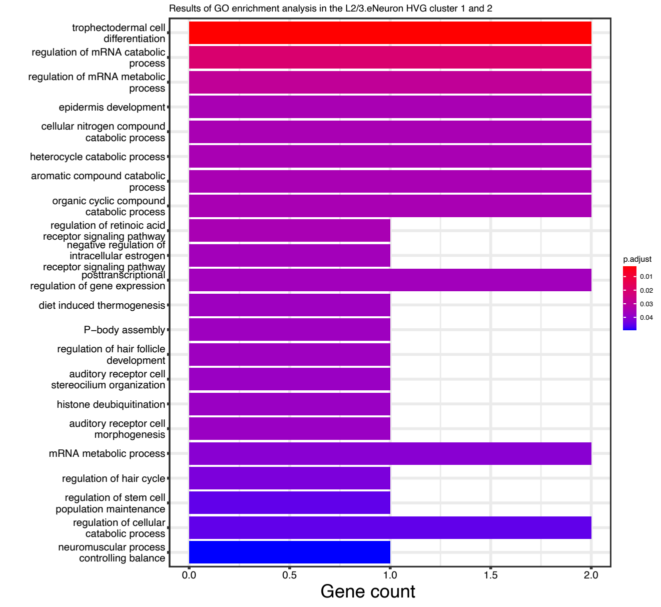


L4 eNeuron

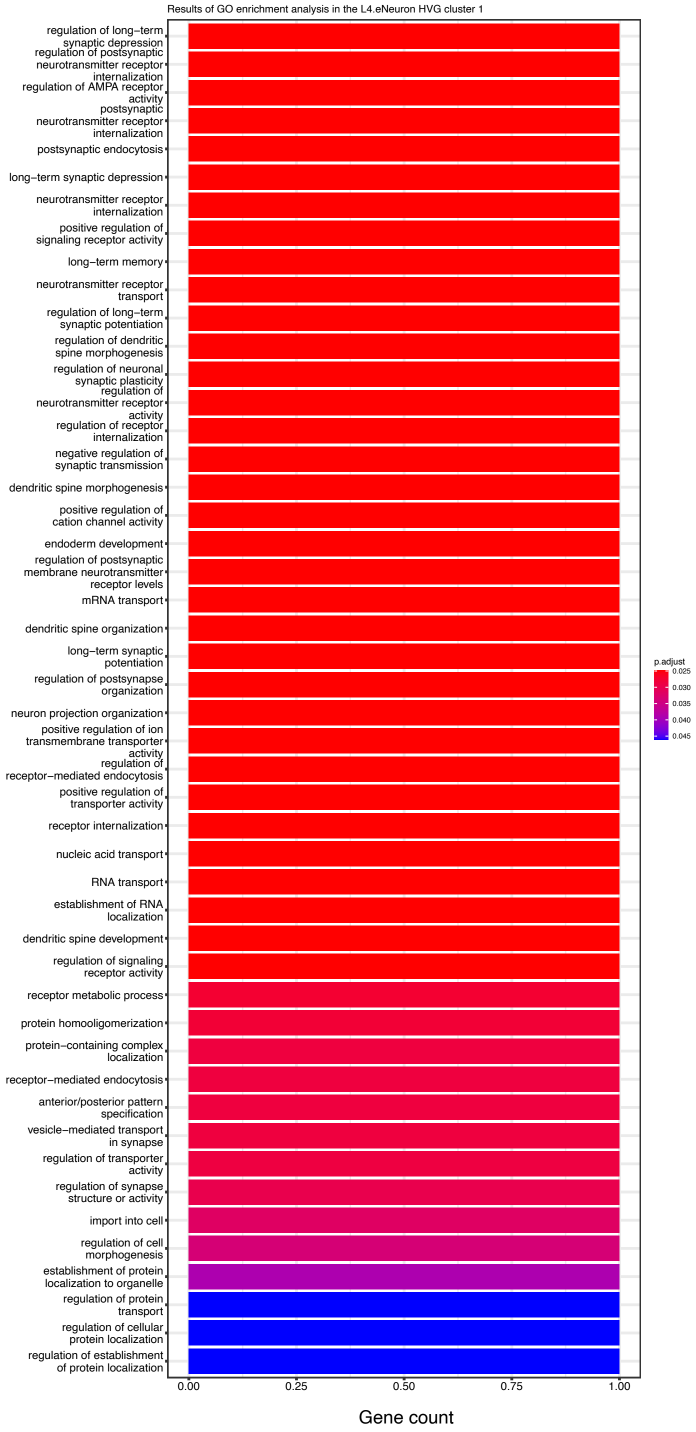


L5 eNeuron

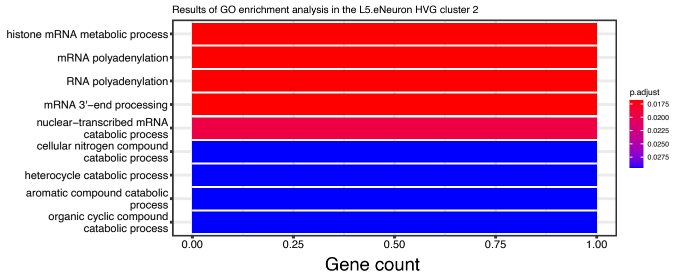


OPCs

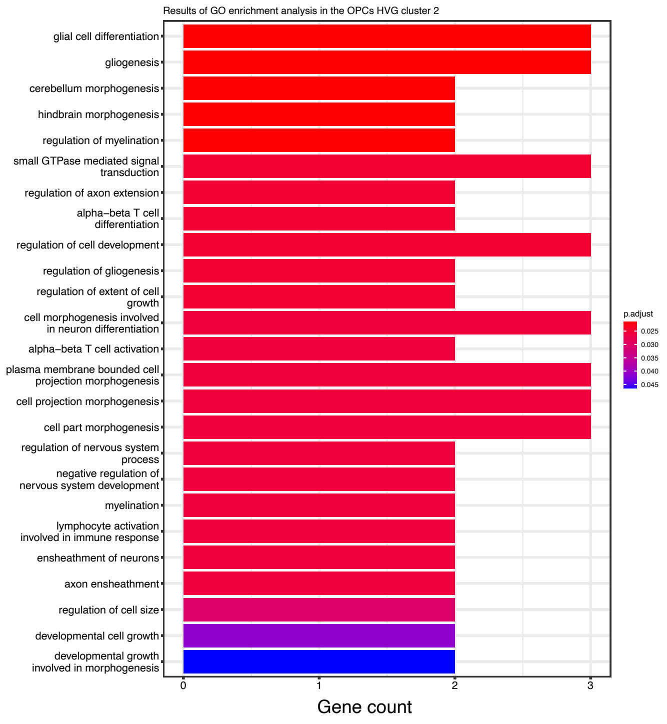


Figure S9. Gene Ontology (GO) enrichment of all the cell types in the seqFISH+ real dataset.

Apllication to the Seq-Scope real dataset:

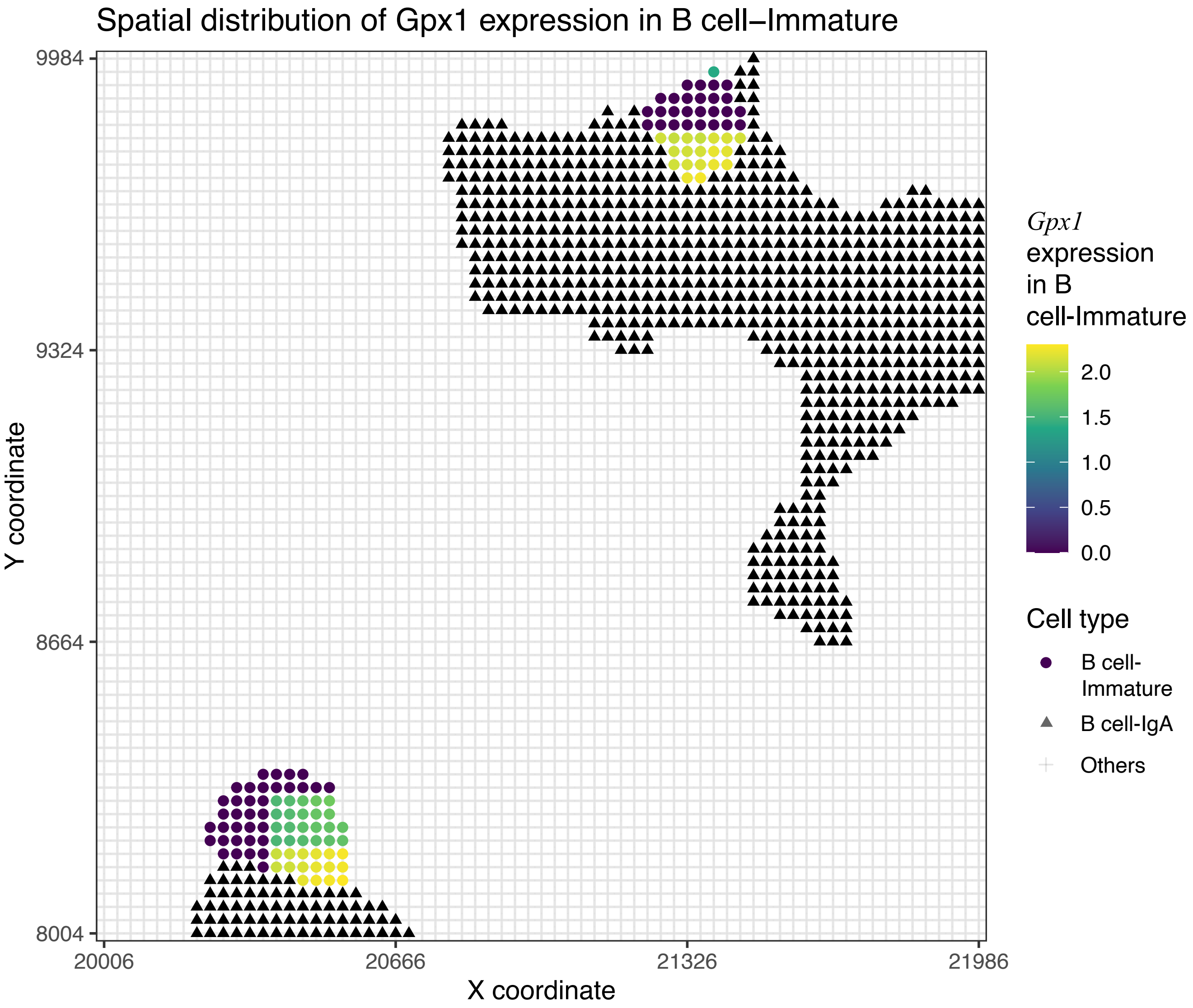


Figure S10. Spatial distribution of *Gpx1* expression in B cell-Immature. The shapes indicate cell types. The color in the circles indicates values of *Gpx1* expression.

### Application to the Seq-Scope real dataset:

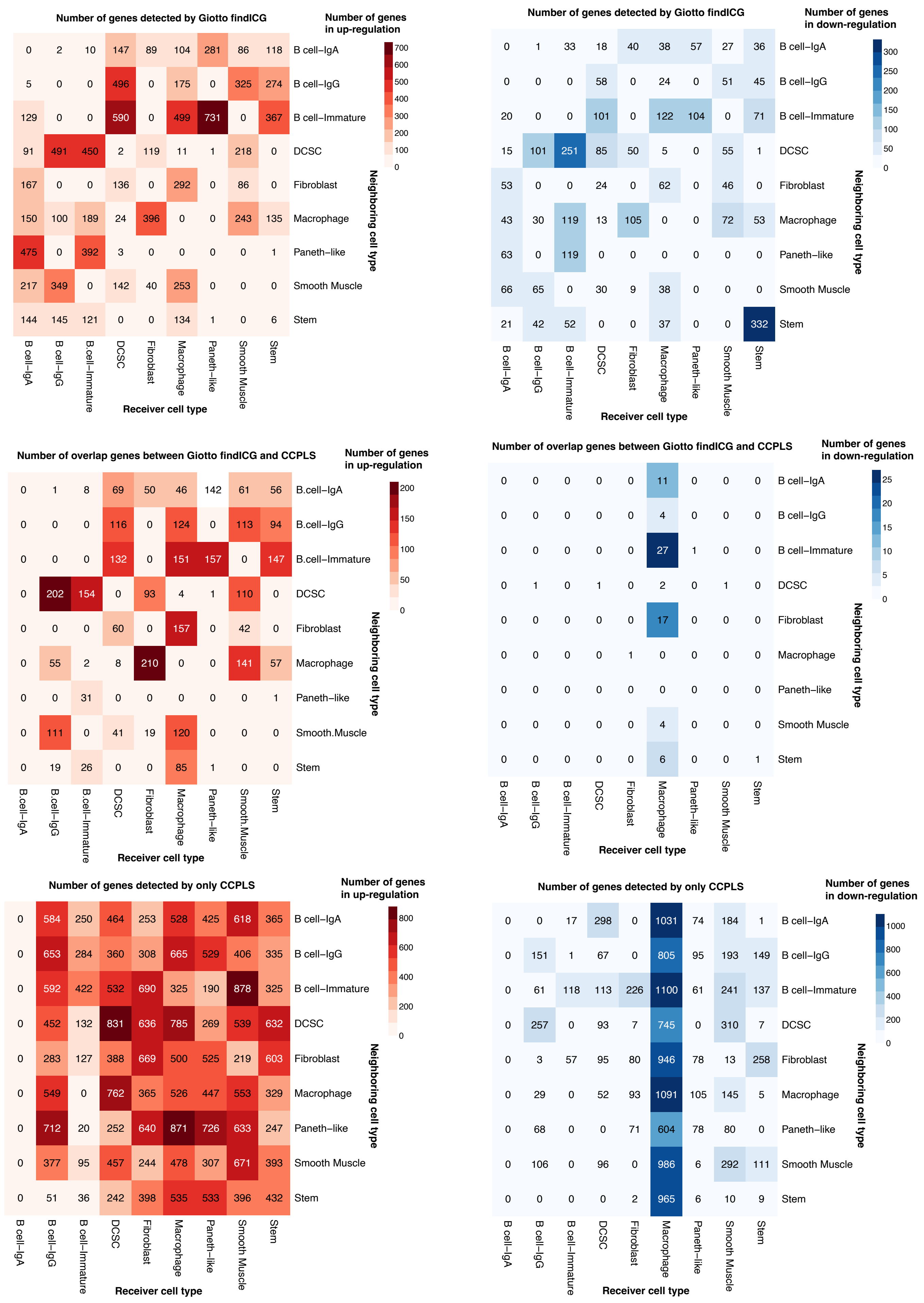


Figure S11. Comparison between Giotto findICG and CCPLS in the Seq-Scope read dataset.

Apllication to the Seq-Scope real dataset:

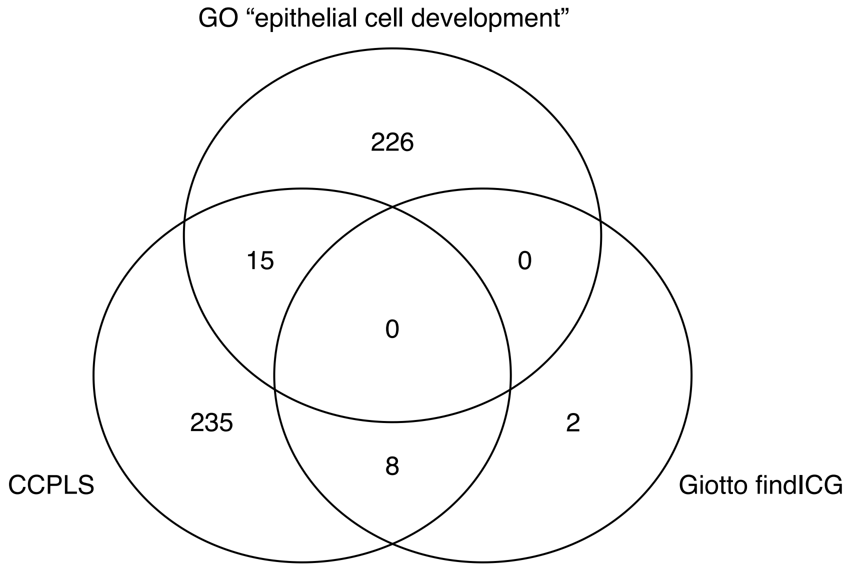


Figure S12. Number of overlaps between genes of GO “epithelial cell development”, genes detected by CCPLS, and genes detected by Giotto findICG. Note that we extracted genes in immature B cell up-regulated by IgA B cell as to CCPLS and Giotto findICG in this venn diagram.

<u>Application to the Seq-Scope real dataset:</u>

Color of heatmap: coefficient

B cell-IgG

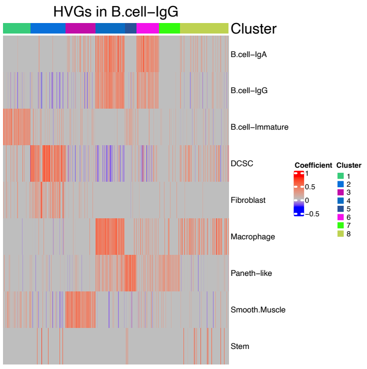


B cell-Immature

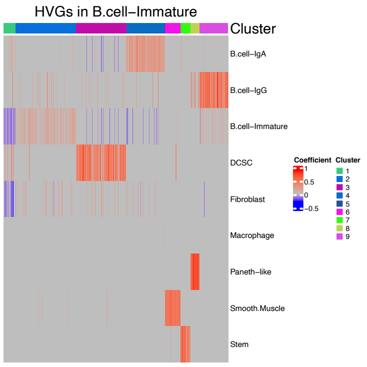


DCSC

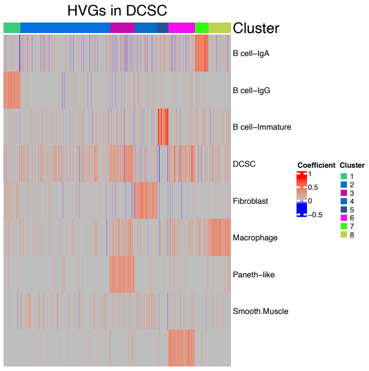


Fibroblast

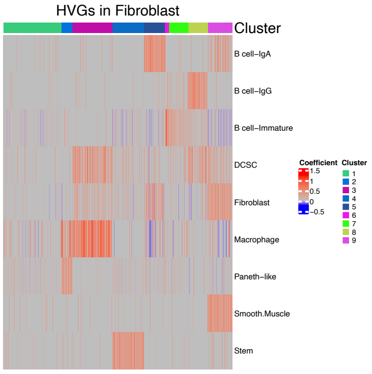


Macrophage

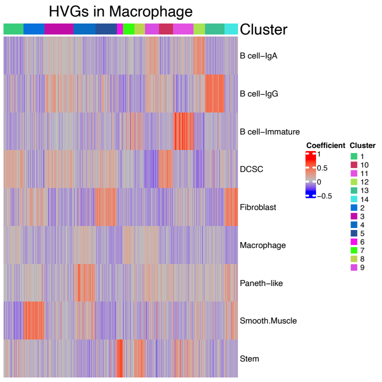


Paneth-like

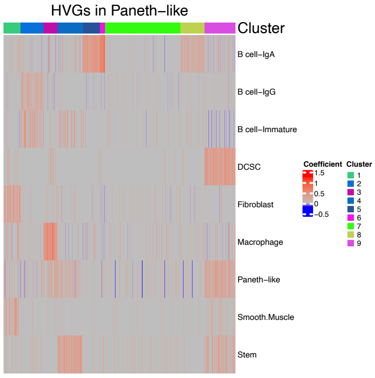


Smooth Muscle

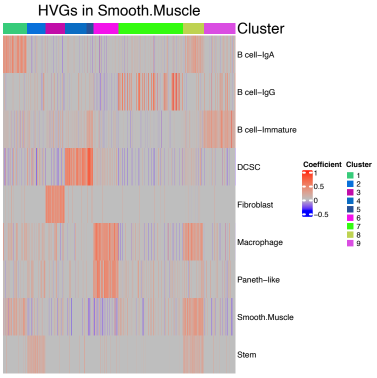


Stem

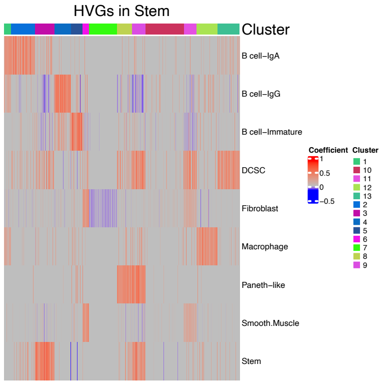


Figure S13. Heat map generated by CCPLS of all the cell types in the Seq-Scope real dataset. Rows and columns correspond to neighboring cell types and highly variable genes (HVGs), respectively. The color of the heat map indicates the coefficient. The heatmap of the immature B cell is the same as that in Figure 4c.

### Application to the Seq-Scope real dataset:

Width of edge: averaged coefficients

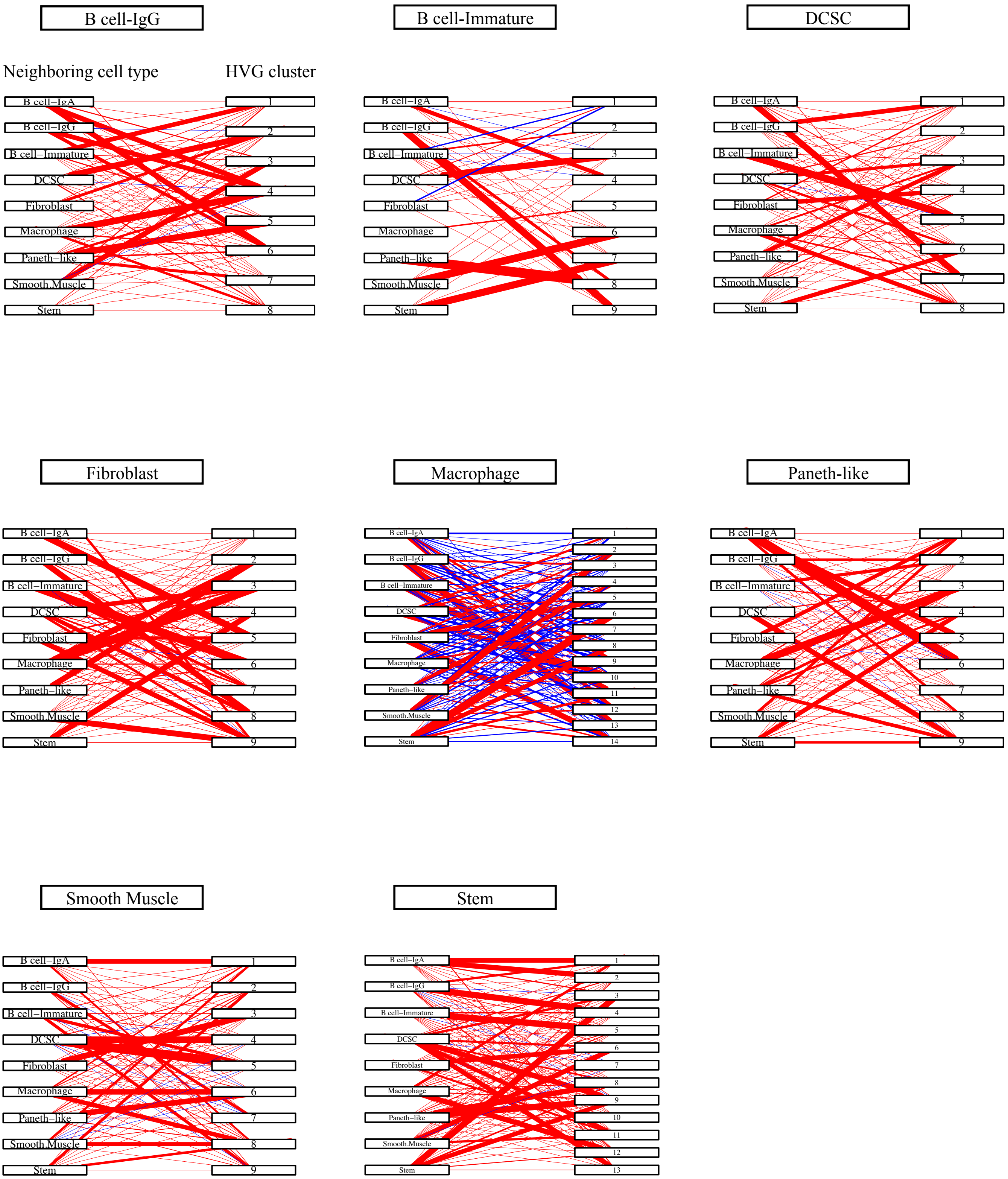


Figure S14. Bipartite graph generated by CCPLS of all the cell types in the Seq-Scope real dataset. The width of each edge indicates the averaged coefficients for each combination of highly variable gene (HVG) clusters and neighboring cell types. The bipartite graph of the immature B cell is the same as that in Figure 4d.

#### Application to the Seq-Scope real dataset:

Red: contributor cell type  Blue: non-contibutor cell typ  Row: neighboring cell type  Column: HVG cluster

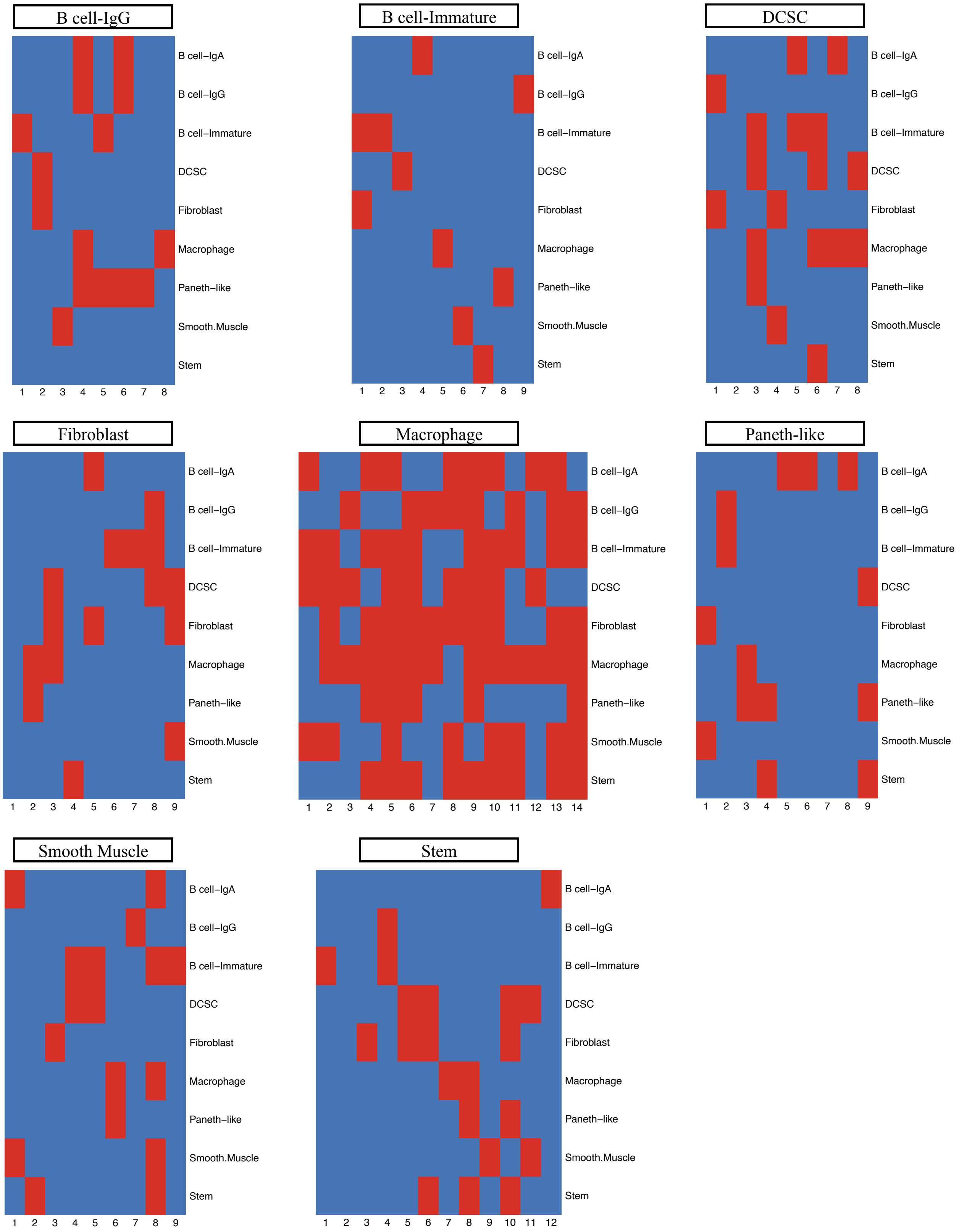


Figure S15. Contributor cell type in the Seq-Scope dataset. The color of the heat map corresponds to the binary value indicating whether the neighbor cell type is a contributor cell type or not. Red and blue indicate the contributor and non-contributor cell types, respectively. Rows and columns correspond to cell types and highly variable gene (HVG) clusters, respectively.

### Application to the Seq-Scope real dataset:

Row: GO term    Column: gene count    Color of bar graph: adjusted *p*-value

B cell-Immature

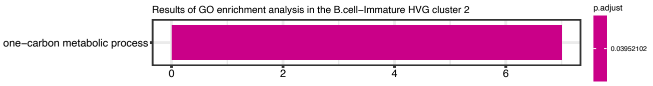


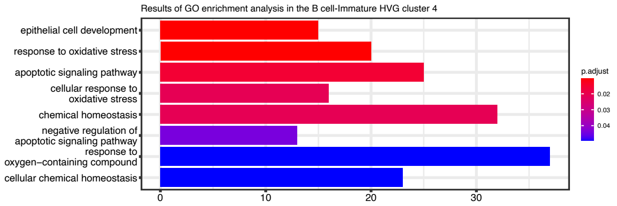


DCSC

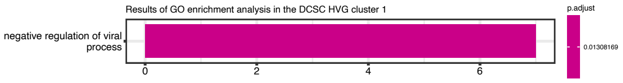


Fibroblast

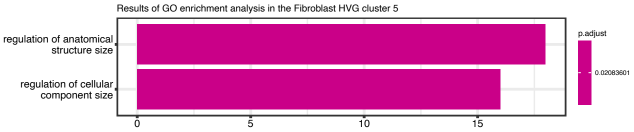


Macrophage

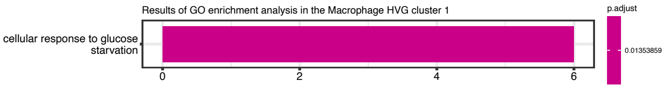


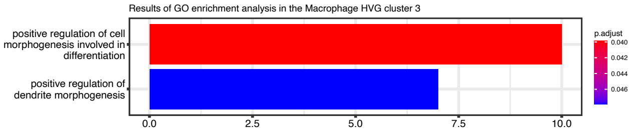


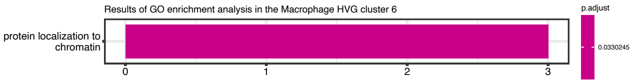


Stem

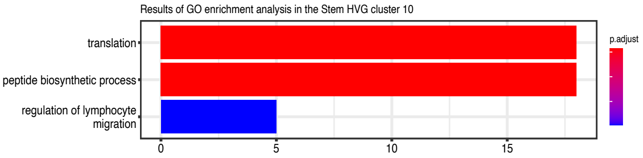


Figure S16. Gene Ontology (GO) enrichment of all the cell types in the Seq-Scope real dataset.
